# Trajectory inference across multiple conditions with condiments: differential topology, progression, differentiation, and expression

**DOI:** 10.1101/2021.03.09.433671

**Authors:** Hector Roux de Bézieux, Koen Van den Berge, Kelly Street, Sandrine Dudoit

## Abstract

In single-cell RNA-sequencing (scRNA-seq), gene expression is assessed individually for each cell, allowing the investigation of developmental processes, such as embryogenesis and cellular differentiation and regeneration, at unprecedented resolutions. In such dynamic biological systems, grouping cells into discrete groups is not reflective of the biology. Cellular states rather form a continuum, e.g., for the differentiation of stem cells into mature cell types. This process is often represented via a trajectory in a reduced-dimensional representation of the scRNA-seq dataset.

While many methods have been suggested for trajectory inference, it is often unclear how to handle multiple biological groups or conditions, e.g., inferring and comparing the differentiation trajectories of wild-type and knock-out stem cell populations.

In this manuscript, we present a method for the estimation and downstream interpretation of cell trajectories across multiple conditions. Our framework allows the interpretation of differences between conditions at the trajectory, cell population, and gene expression levels. We start by integrating datasets from multiple conditions into a single trajectory. By comparing the conditions along the trajectory’s path, we can detect large-scale changes, indicative of differential progression. We also demonstrate how to detect subtler changes by finding genes that exhibit different behaviors between these conditions along a differentiation path.

---

The emergence of RNA sequencing at the single-cell level (scRNA-Seq) has enabled a new degree of resolution in the study of cellular processes. The ability to consider biological processes as continuous phenomena instead of individual discrete stages has permitted a finer and more comprehensive understanding of dynamic processes such as embryogenesis and cellular differentiation. Trajectory inference was one of the first applications that leveraged this continuum [1] and a consequential number of methods have been proposed since then [2–4]. Saelens et al. [5] offer an extensive overview and comparison of such methods. Analysis of scRNA-Seq datasets using a curated database reveals that about half of all datasets were used for trajectory inference (TI) [6]. At its core, TI represents a dynamic process as a directed graph. Distinct paths along this graph are called lineages. Individual cells are then projected onto these lineages and the distance along each path is called pseudotime. In this setting, developmental processes are often represented in a tree structure, while cell cycles are represented as a loop. Following TI, other methods have been proposed to investigate differential expression (DE) along or between lineages, either as parts of TI methods [3, 7] or as separate modules that can be combined to create a full pipeline [8].

More recently, other methods have emerged to answer an orthogonal problem, focusing on systems under multiple conditions. This includes, for example, situations where a biological process is studied both under a normal (or control) condition and under an intervention such as a treatment [9–11] or a genetic modification [12]. In other instances, one may want to contrast healthy versus diseased [13] cells or even more than two conditions [14]. In such settings, one might look for differential abundance, i.e., cell population shifts between conditions. Initial analytical approaches ignored the continuous nature of biological processes and binned cells into discrete clusters before looking at differences in composition between clusters. Borrowing from the field of mass cytometry [15], `milo` [16], and `DAseq` [17] rely on low-dimensional representations of the observations and define data-driven local neighborhoods in which they test for differences in compositions. Each of these methods show clear improvements in performance over cluster-based methods, and provide a more principled approach that better reflects the nature of the system.

However, many studies with multiple conditions, if not most, actually involve processes that can be described by a trajectory. Utilizing this underlying biology could increase either the interpretability of the results or the ability to detect true and meaningful changes between conditions. In this manuscript, we present the `condiments` workflow, a general framework to analyze dynamic processes under multiple conditions that leverages the concept of a trajectory structure. `condiments` has a more specific focus than `milo` or `DAseq`, but it compensates for this by improving the quality of the differential abundance assessment and its biological interpretation. Our proposed analysis workflow is divided into three steps. In Step 1, `condiments` considers the trajectory inference question, assessing whether the dynamic process is fundamentally different between conditions, which we call *differential topology.* In Step 2, it tests for differential abundance of the different conditions along lineages and between lineages, which we respectively call *differential progression* and *differential differentiation.* Lastly, in Step 3, it estimates gene expression profiles similarly to Van den Berge et al. [8] and tests whether gene expression patterns differ between conditions along lineages, therefore extending the scope of `differential expression`.

In this manuscript, we first present the `condiments` workflow, by detailing the underlying statistical model, and providing an explanation and intuition for each step. We then benchmark `condiments` against more general methods that test for differential abundance to showcase how leveraging the existence of a trajectory improves the assessment of differential abundance. Finally, we demonstrate the flexibility and improved interpretability of the `condiments` workflow in three case studies that span a variety of biological settings and topologies.

## Results

### General model and workflow

#### Data structure and statistical model

We observe gene expression measures for *J* genes in *n* cells, resulting in a *J × n* count matrix **Y**. For each cell *i*, we also know its condition label *c*(*i*) ∈ {1,…, *C*} (e.g.,”treatment” or “control”, “knock-out” or “wild-type”). We assume that, for each condition *c*, there is an underlying developmental structure 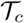, or trajectory, that possesses a set of *L_c_* lineages.

For a given cell i with condition *c*(*i*), its position along the developmental path 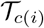 is defined by a vector of *L*_*c*(*i*)_ pseudotimes **T**_*i*_ and a unit-norm vector of *L*_*c*(*i*)_ weights **W**_*i*_ (||**W**_*i*_||_1_ = 1) (i.e., there is one pseudotime and one weight per lineage), with

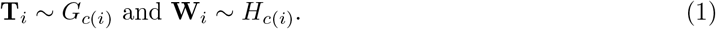

The cumulative distribution functions (CDF) *G_c_* and *H_c_* are condition-specific and we make limited assumptions on their properties (see the method section for details). The pseudotime values represent how far a cell has progressed along each lineage, while the weights represent how likely it is that a cell belongs to each lineage. The gene expression model will be described below. Using this notation, we can properly define a trajectory inference (TI) method as a function that takes as input **Y** – and potentially other arguments – and returns estimates of *L_c_*, **T**, **W**, and eventually 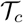.

#### Step 1 - Differential Topology: Should we fit a common trajectory?

The first question to ask in our workflow is: Should we fit a common trajectory to all cells regardless of their condition? Or are the developmental trajectories too dissimilar between conditions? To demonstrate what this means, consider two extremes. For a dataset that consists of a mix of bone marrow stem cells and epithelial stem cells, using tissue as our condition, it is obvious that the developmental trajectories of the two conditions are not identical and should be estimated separately. On the other hand, if we consider a dataset where only a few genes are differentially expressed between conditions, the impact on the developmental process will be minimal and it is sensible to estimate a single common trajectory.

Indeed, we favor fitting a common trajectory for several reasons. Firstly, fitting a common trajectory is a more stable procedure since more cells are used to infer the trajectory. Secondly, our workflow still provides a way to test for differences between conditions along and between lineages even if a common trajectory is inferred. In particular, fitting a common trajectory between conditions does not require that cells of distinct conditions differentiate similarly along that trajectory. Finally, fitting different trajectories greatly complicates downstream analyses since we may need to map between distinct developmental structures before comparing them (i.e., each lineage in the first trajectory must match exactly one lineage in the second trajectory). Therefore, our workflow recommends fitting a common trajectory if the differences between conditions are *small enough.*

To quantify what *small enough* is, we rely on two approaches. The first is a qualitative diagnostic tool called *imbalance score*. It requires as input a reduced-dimensional representation **X** of the data **Y** and the condition labels. Each cell is assigned a score that measures the imbalance between the local and global distributions of condition labels. Similarly to Dann et al. [16], Burkhardt et al. [18], the neighborhood of a cell is defined using a *k*-nearest neighbor graph on **X**, which allows the method to scale very well to large values of *n*. Cell-level scores are then locally scaled using smoothers in the reduced-dimensional space (see the Methods section).

However, visual representation of the scores may not always be enough to decide whether or not to fit a common trajectory in less obvious cases. Therefore, we introduce a more principled approach, the `topologyTest`. This test assesses whether we can reject the following null hypothesis:

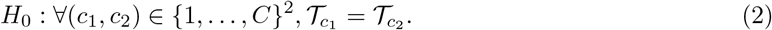

Under the null, the trajectory is common among all conditions and can therefore be estimated using all cells. Therefore, an estimation of the pseudotime vectors done by inferring a trajectory for each condition should be equivalent to the same procedure after permuting the condition labels. This is what is done for the `topologyTest`. A set of pseudotime vectors is estimated with the true condition labels. Another set is generated using permuted labels. Under the null, these two distributions should be equal. We can therefore test hypothesis (2) by testing for the equality in distributions of pseudotime using a variety of statistical tests (see the Methods section for details). Since we want to favor fitting a common trajectory and we only want to discover cases that are not only statistically significant but also biologically relevant, the tests typically include a minimum magnitude requirement for considering the difference between distributions to be significant (similar to a minimum log-fold-change for assessing DE). More details and practical implementation considerations are discussed in the Methods section.

In practice, the `topologyTest` requires maintaining a mapping between each of the trajectories, both between conditions and between permutations (see the Methods section where we define a mapping precisely). Trajectory inference remains a semi-supervised task, that generally cannot be fully automated. In particular, the number of estimated lineages might change between different permutations for a given condition, precluding a mapping. As such, the `topologyTest` is only compatible with certain TI methods that allow for the specification of an underlying skeleton structure [2, 4], where the adjacency matrix can be pre-specified, as well (optionally) start and/or end states.

In the examples from Fig 1, the skeleton of the trajectory is represented by a series of nodes and edges. In examples 1b-d, the knock-out has no impact on this skeleton compared to the wild-type. In example 1e, the knock-out (KO) modifies the skeleton, in that the locations of the nodes change. However, the adjacency matrix does not change and the two skeletons represent isomorphic graphs: the skeleton structure is preserved.

**Figure 1:**
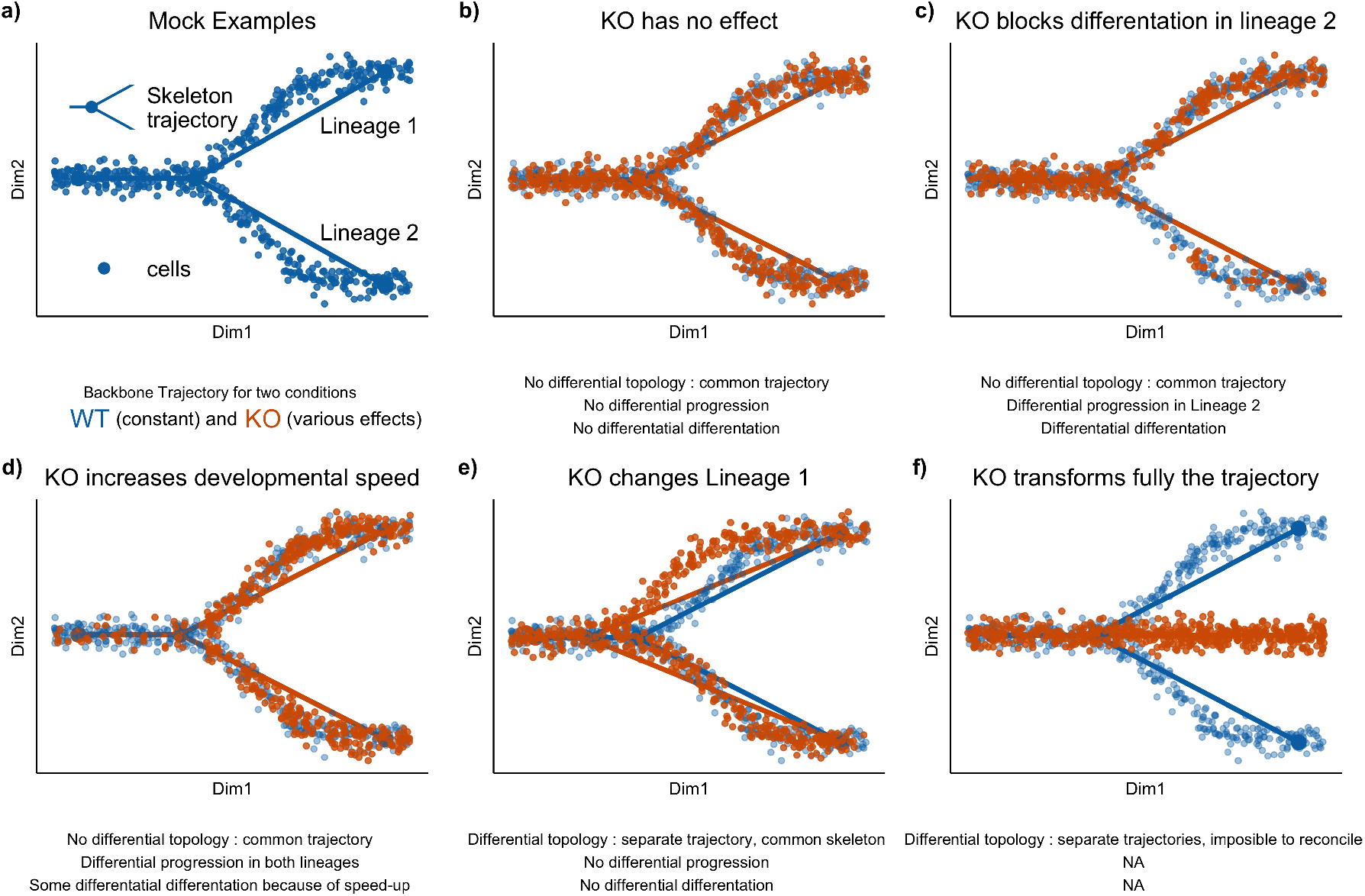
Illustrating the first two steps of the condiments workflow with several scenarios. **(a.)** The examples are all built on a similar wild-type backbone, i.e., two lineages that slowly diverge in the absence of knocking out. Cells either originate from a wild-type (WT, blue) or a knock-out (KO, orange) condition. In **(b.)**, the knock-out has no effect, all three tests fail to reject their null hypothesis. In **(c.)**, the knock-out partly blocks differentiation along Lineage 2, meaning that fewer cells develop along that lineage. In this case, while the `topologyTest` fails to reject the null, we have both differential progression along Lineage 2 and differential differentiation. In **(d.)**, the knock-out speeds development, so there are more orange cells toward the end of both lineages. This leads to both differential progression and differentiation. In **(e.)**, the knock-out modifies the intermediate stage for Lineage 1 and changes where the lineages bifurcates; based on the `topologyTest`, we fit one trajectory per condition. However, the skeleton structure is unchanged, so there is a mapping between the two trajectories and we can still test for differential progression and differentiation. In both cases, we fail to reject the null. Finally, in **(f.)**, the knock-out fully disrupts the developmental process: all cells in the knock-out condition progress along a new lineage. Here, we fit separate trajectories and these cannot be reconciled easily, so we cannot proceed to Steps 2 and 3.

For some TI methods [2, 4], it is possible to specify and preserve this skeleton structure. This means that the mapping of lineages can be done automatically. The `topologyTest` utilises this, and is thus restricted to such TI methods. This common skeleton structure can also be used if the null of the `topologyTest` is rejected. The availability of a mapping between lineages means that the next steps of the workflow can be conducted as if we had failed to reject the null hypothesis, as done in Fig 1e. The third case study will also present an example of this.

Even if the null is rejected by the `topologyTest` and separate trajectories must be fitted for each condition, a common skeleton structure can still be used to map between trajectories. This mapping means that the next steps of the workflow can be conducted as if we had failed to reject the null hypothesis, as done in Fig 1e. The third case study will also present an example of this. In cases where no common skeleton structure exists, such as Fig 1f, no automatic mapping exists. Differential abundance can be assessed but requires a manual mapping. Differential expression can still be conducted as well.

#### Step 2 - Differential abundance: What are the global differences between conditions?

The second step of the workflow focuses on differences between conditions at the trajectory level. It requires either a common trajectory, or multiple trajectories and a mapping. We can then ask whether cells from different conditions behave similarly as they progress along the trajectory. To facilitate the interpretation of the results, we break this into two separate questions. Note that, at this step and the next, we are no longer limited to specific TI methods. Moreover, the mapping can be partial. In that case, Step 2 will be restricted to the parts (or subgraphs) of the trajectories that are mappable. See the Methods section for proper definitions of mapping and partial mapping.

#### Step 2a: Differential Progression

Although the topology might be common, cells might progress at different rates along the lineages for different conditions. For example, a treatment might limit the differentiation potential of the cells compared to the control, or instead speed it up. In the first case, one would expect to have more cells at the early stages and fewer at the terminal state, when comparing treatment and control. Using our statistical framework, testing for differential progression amounts to testing:

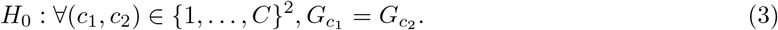

This test can also be conducted at the individual-lineage level. If we denote by *G_lc_* the *l^th^* component of the distribution function *G_c_*, we can test for differential progression along lineage *l* by considering the null hypothesis:

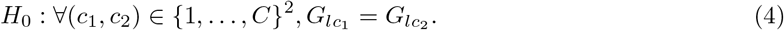

We can assess either or both null hypotheses in the `progressionTest`, which relies on non-parametric tests to compare two or more distributions, e.g., the Kolmogorov-Smirnov test [19] or the classifier test [20]. More details and practical implementation considerations are discussed in the Methods Section.

#### Step 2b: Differential Differentiation

Although the topology might be common, cells might also differentiate in varying proportions between the lineages for different conditions. For example, an intervention might lead to preferential differentiation along one lineage over another, compared to the control condition; or might alter survival rates of differentiated cells between two end states. In both cases, the weight distribution will be different between the control and treatment. Assessing differential differentiation at the global level amounts to testing, in our statistical framework, the null hypothesis

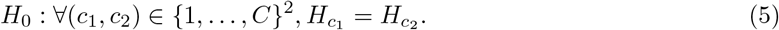

This test can also be conducted for a single pair of lineages (*l, l*’):

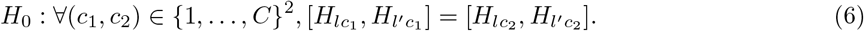

The above null hypotheses can again be tested by relying on non-parametric test statistics. We also discuss specific details and practical implementation in the Methods section.

The `progressionTest` and `differentiationTest` are quite linked since the functions *G_c_* and *H_c_* are correlated and will therefore often return similar results. However, they do answer somewhat different questions. In particular, looking at single-lineage (`progressionTest`) and lineage-pair (`differentiationTest`) test statistics will allow for a better understanding of the global differences between conditions. Differential differentiation does not necessarily imply differential progression and vice versa.

#### Step 3 - Differential Expression: Which genes have different expression patterns between conditions?

Steps 1 and 2 focus on differences at a global level (i.e., aggregated over all genes) and will detect large changes between conditions. However, such major changes are ultimately driven by underlying differences in gene expression patterns. Furthermore, even in the absence of global differences, conditions might still have some more subtle impact at the gene level. In the third step, we therefore compare gene expression patterns between conditions for each of the lineages. Step 3 is even more general than Step 2, in that it can be used without mapping between trajectories, i.e., some or all lineages could be condition-specific.

Following the `tradeSeq` manuscript by Van den Berge et al. [8], we consider a general and flexible model for gene expression, where the gene expression measure *Y_ji_* for gene *j* in cell *i* is modeled with the negative binomial generalized additive model (NB-GAM) described in Equation (13). We extend the `tradeSeq` model by additionally estimating condition-specific average gene expression profiles for each gene. We therefore rely on lineage-specific, gene-specific, and condition-specific smoothers, *s_jlc_*. With this notation, we can introduce the `conditionTest`, which, for a given gene *j*, tests the null hypothesis that these smoothers are identical across conditions:

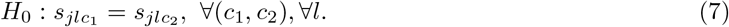

As in `tradeSeq`, we rely on the Wald test to test *H*_0_ in terms of the smoothers’ regression coefficients. We can also use the fitted smoothers to visualize gene expression along lineages between conditions or cluster genes according to their expression patterns.

### Simulations

We generate multiple trajectories using the simulation framework provided by Cannoodt et al. [22]. Within this framework, it is possible to knock out a specific gene. Here, we knock out a master regulator that drives differentiation into the second lineage. The strength of this knock-out can be controlled via a multiplier parameter *m*. If *m* = 0, the knock-out is total. If 0 < *m* < 1, we have partial knock-out. If *m* > 1, the master regulator is over-expressed and cells differentiate much faster along the second lineage.

Three types of datasets are generated: Simple branching trajectories (two lineages, e.g., Fig. 2a) of 3, 500 cells, with equal parts wild-type and knock-out; trajectories with two consecutive branchings (and thus three lineages, e.g., Fig. 2b) of 3, 500 cells, with equal parts wild-type and knock-out; and branching trajectories (two lineages) of 5, 000 cells with three conditions, wild-type, knock-out with multiplier *m*, and induction with multiplier 1/*m* (Fig. 2c).

**Figure 2:**
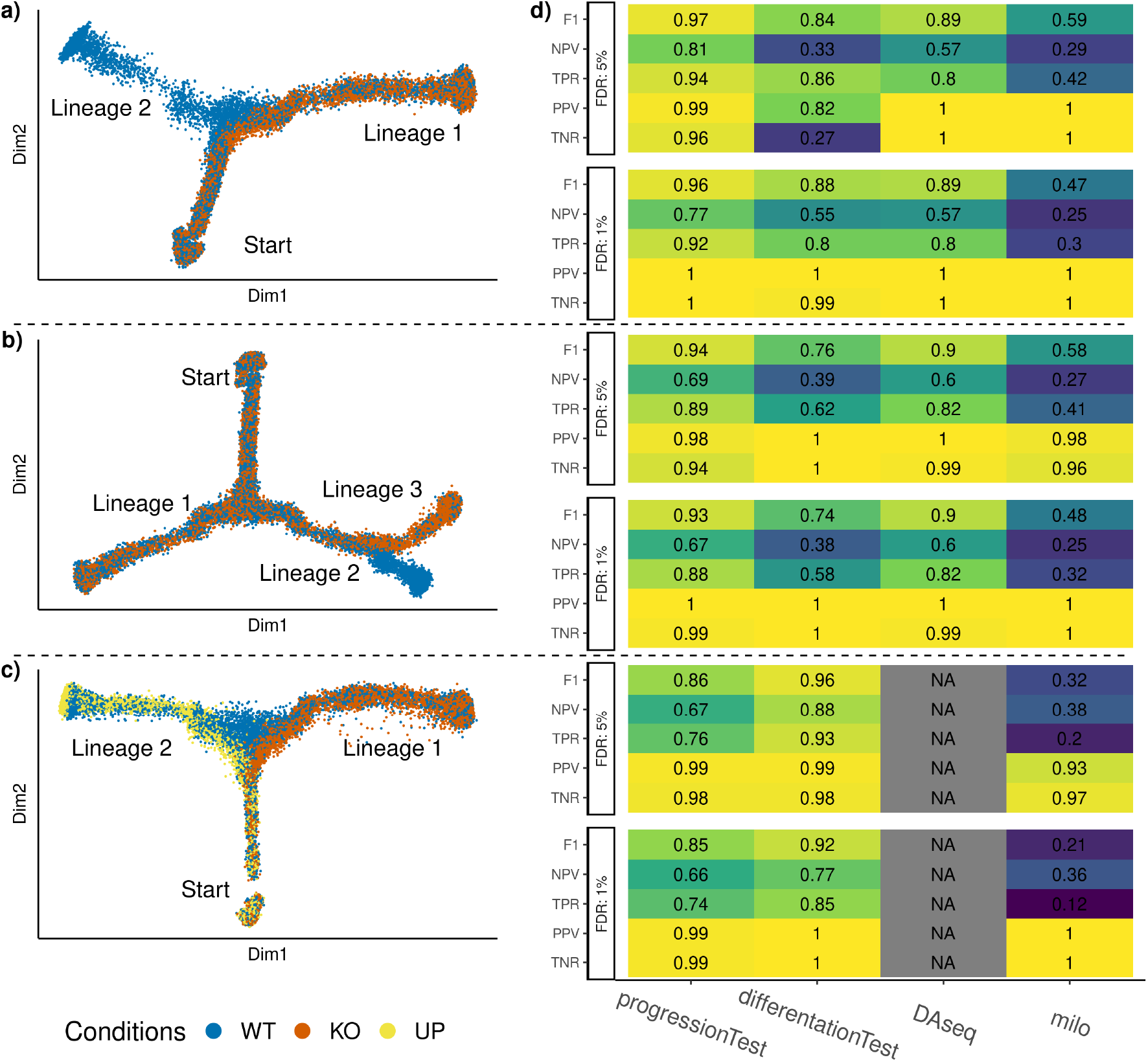
Simulation results. Three types of datasets are generated, with respectively two, three, and two lineages, and two, two, and three conditions. Reduced-dimensional representations of these datasets, for a multiplier value of *m* = .5, are presented in **(a.)**, **(b.)**, and **(c.)**, respectively. After generating multiple versions of the datasets for a range of values of m, we compare the performance of the `progressionTest` and `differentiationTest` with that of `DAseq`[17] and `milo`[16], when controlling the false discovery rate at nominal levels 1% and 5% using the Benjamini-Hochberg [21] procedure. In **(d.)**, each cell represents the performance measure associated with one test on one dataset for one nominal FDR level. Cells are also colored according to the performance. Overall, with two conditions, the progressionTest ranks first, followed by `DAseq` and the `differentiationTest`. With three conditions, the `differentiationTest` ranks first. `DAseq` is limited to two conditions. Exact simulation parameters and metrics are specified in the Methods section.

The simulation framework cannot, however, generate distinct trajectories for the different conditions, so we start the `condiments` workflow at Step 2, downstream of `slingshot`. We compare the `progressionTest` and `differentiationTest` from `condiments` to methods that also do not rely on clustering, but instead take into account the continuum of differentiation. `milo` [16] and `DAseq`[17] both define local neighborhoods using *k*-nearest neighbors graphs and look at differences of proportions in these neighborhood to test for differential abundance. These methods returns multiple tests per dataset (i.e., one per neighborhood), so we adjustfor multiple hypothesis testing using the Benjamini-Hochberg procedure [21]. By applying `milo`, `DAseq`, and `condiments` on the simulated datasets, we can compare the results of the tests versus the values of *m*: We count a true positive when a test rejects the null and *m* ≠ 1, and a true negative if the test fails to reject the null and *m* = 1.

We compare the methods’ ability to detect correct differences between conditions using five metrics: The true negative rate (TNR), positive predictive value (PPV), true positive rate (TPR), negative predictive value (NPV), and F1-score, when controlling the FDR at two nominal levels of 1% and 5%. More details on the simulation scenarios and metrics can be found in the Methods section. Results are displayed in Fig. 2d.

On all simulations, all methods display excellent results for the TNR and PPV (except for the `differentiationTest` with level 1% on the branching dataset). However, the performances for the TPR (power), NPV, and F1-rate vary quite widely. On the two types of datasets with two conditions, the ranking is uniform over all metrics and levels: `progressionTest, DAseq, differentiationTest`, and `milo`. On the third simulation setting with three conditions, we cannot benchmark `DAseq` since its testing framework is restricted to two conditions. Here, also, the ranking is uniform but the `differentiationTest` outperforms the `progressionTest`. Looking more closely at the results, we can see (Fig S2) that this mostly stems from increased power for the `differentiationTest` when *m* is close to 1.

Overall, the tests from the `condiments` workflow offer a flexible approach that can handle various scenarios and still outperform competitors.

### Case studies

We consider three real datasets as case studies for the application of the `condiments` workflow. Table 1 gives an overview of these datasets and summary results. These case studies aim to demonstrate the versatility and usefulness of the `condiments` workflow, as well as showcase how to interpret and use the tests in practice.

**Table 1:** Summary of all case study datasets. We report the name, number of cells *n,* number of conditions *C*, number of lineages *L* of each dataset, as well as the p-value resulting from testing for differential topology, progression and differentiation and the number of differentially expressed genes between conditions according to the conditionTest

| Dataset | $n$ | $C$ | $L$ | topology | progression | differentiation | DE |
| --- | --- | --- | --- | --- | --- | --- | --- |
| **TGFB**[9] | 9,268 | 2 | 1 | 0.38 | $\leq 2.2 \times 10^{-16}$ | NA | 1,993 |
| **TCDD**[10] | 9,951 | 2 | 1 | 0.07 | $\leq 2.2 \times 10^{-16}$ | NA | 2,144 |
| **KRAS**[23] | 10,177 | 3 | 3 | $\leq 2.2 \times 10^{-16}$ | $\leq 2.2 \times 10^{-16}$ | $\leq 2.2 \times 10^{-16}$ | 363 |

### TGFB dataset

McFaline-Figueroa et al. [9] studied the epithelial-to-mesenchymal transition (EMT), where cells migrate from the epithelium (inner part of the tissue culture dish) to the mesenchyme (outer part of the tissue culture dish) during development. The developmental process therefore is both temporal and spatial. As cells differentiate, gene expression changes. Moreover, the authors studied this system under two settings: a mock (control) condition and a condition under activation of transforming growth factor *β* (TGFB).

After pre-processing, normalization, and integration (see details in the supplementary methods), we have a dataset of 9, 268 cells, of which 5, 207 are mock and 4, 241 are TGFB-activated. The dataset is represented in reduced dimension using UMAP[24] (Fig. 3a). Adding the spatial label of the cells (Fig. 3b) shows that the reduced-dimensional representation of the gene expression data captures the differentiation process.

**Figure 3:**
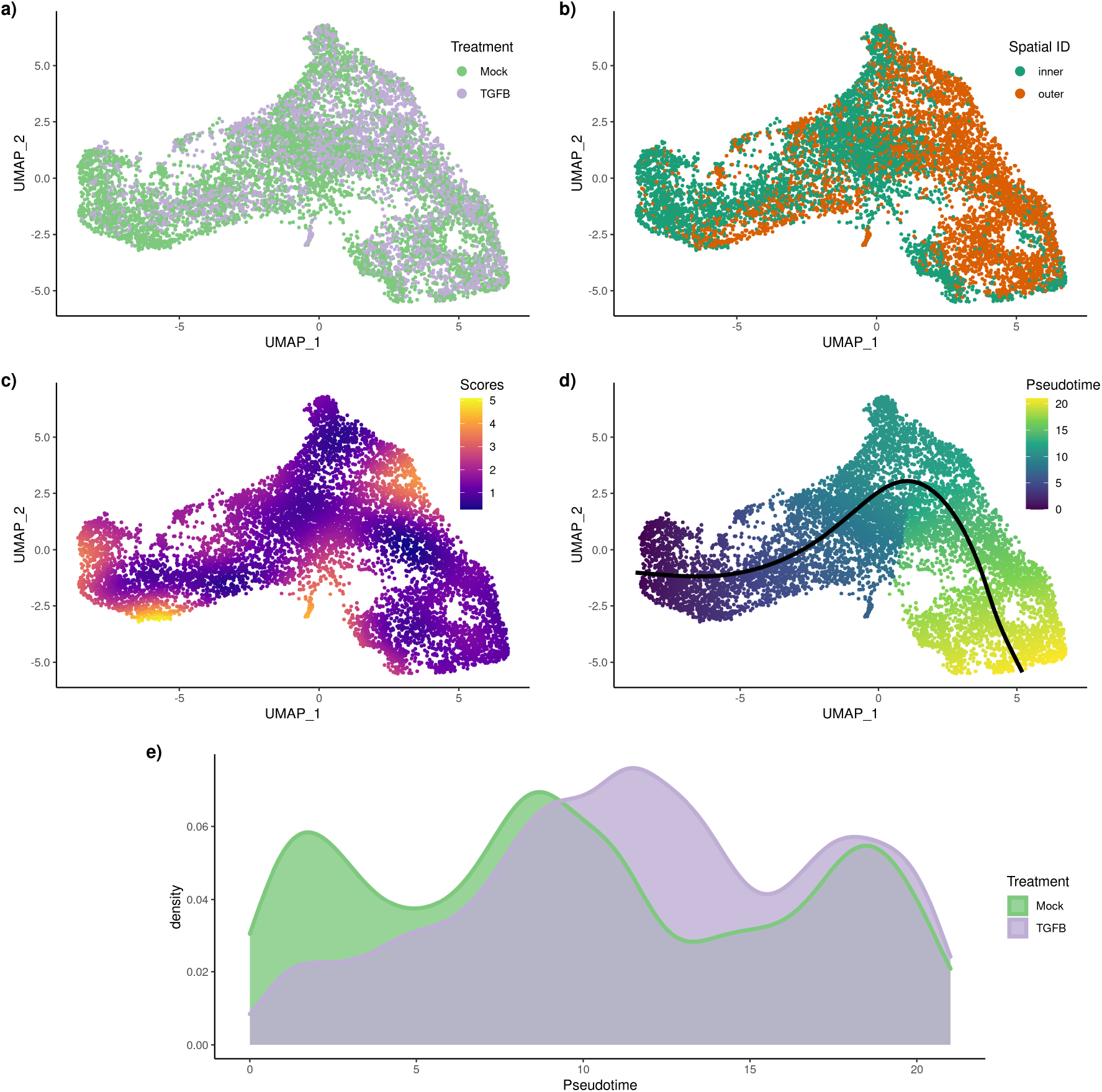
**TGFB** dataset: Differential topology and differential progression. After normalization and projection on a reduced-dimensional space (using UMAP), the cells can be colored either by treatment label **(a.)** or spatial origin **(b.)**. Using the treatment label and the reduced-dimensional coordinates, an imbalance score is computed and displayed **(c.)**. The `topologyTest` fails to reject the null hypothesis of no differential topology and a common trajectory is therefore fitted **(d.)**. However, there is differential progression between conditions: the pseudotime distributions along the trajectory are not identical **(e.)** between conditions and we reject the null using the `progressionTest`.

We can then run the `condiments` workflow. The imbalance score of each cell is computed and displayed in Fig. 3c. Although some regions do display strong imbalance, there is no specific pattern along the developmental path. This is confirmed when we run the `topologyTest`. The nominal *p*-value of the associated test is 0.38. We clearly fail to reject the null hypothesis and we consequently fit a common trajectory to both conditions using `slingshot` with the spatial labels as clusters. This single-lineage trajectory is shown in Fig. 3d.

Next, we can ask whether the TGFB treatment impacts the differentiation speed. The developmental stage of each cell is estimated using its pseudotime. Plotting the per-condition kernel density estimates of pseudotimes in Fig. 3e reveals a strong treatment effect. The pseudotime distribution for the mock cells is trimodal, likely reflecting initial, intermediary, and terminal states. However, the first mode is not present in the TGFB condition, and the second is skewed towards higher pseudotime values. This is very consistent with the fact that the treatment is a growth factor that would increase differentiation, as shown in the original publication. Testing for equality of the two distributions with the `progressionTest` confirms the visual interpretation. The nominal *p*-value associated with the test is smaller than 2.2 × 10^-16^ and we reject the null that the distributions are identical. Since the trajectory is limited to one lineage, there is no possible differential differentiation between pairs of lineages.

Then, we proceed to identifying genes whose expression patterns differ between the mock and TGFB conditions. After gene filtering, we fit smoothers to 10, 549 genes, relying on the model described in Equation (13). We test whether the smoothers are significantly different between conditions using the `conditionTest`. Testing against a log-fold-change threshold of 2, we find 1, 993 genes that are dynamically differentially expressed between the two conditions when controlling the false discovery rate (FDR) at a nominal level of 5%. Fig. 4a and b show the two genes with the highest Wald test statistic. The first gene, *LAMC2*, was also found to be differentially expressed in the original publication and has been shown to regulate EMT [25]. The second gene, *TGFBI* or TGFB-induced gene, is not surprising, and was also labelled as differentially expressed in the original publication. In contrast, the gene that is deemed the least differentially expressed exhibits no differences between the smoothers (Fig. 4c.). Looking at all 1, 993 DE genes, we can cluster and display their expression patterns along the lineage for both conditions (Fig. 4) and identify several groups of genes that have different patterns between the two conditions.

**Figure 4:**
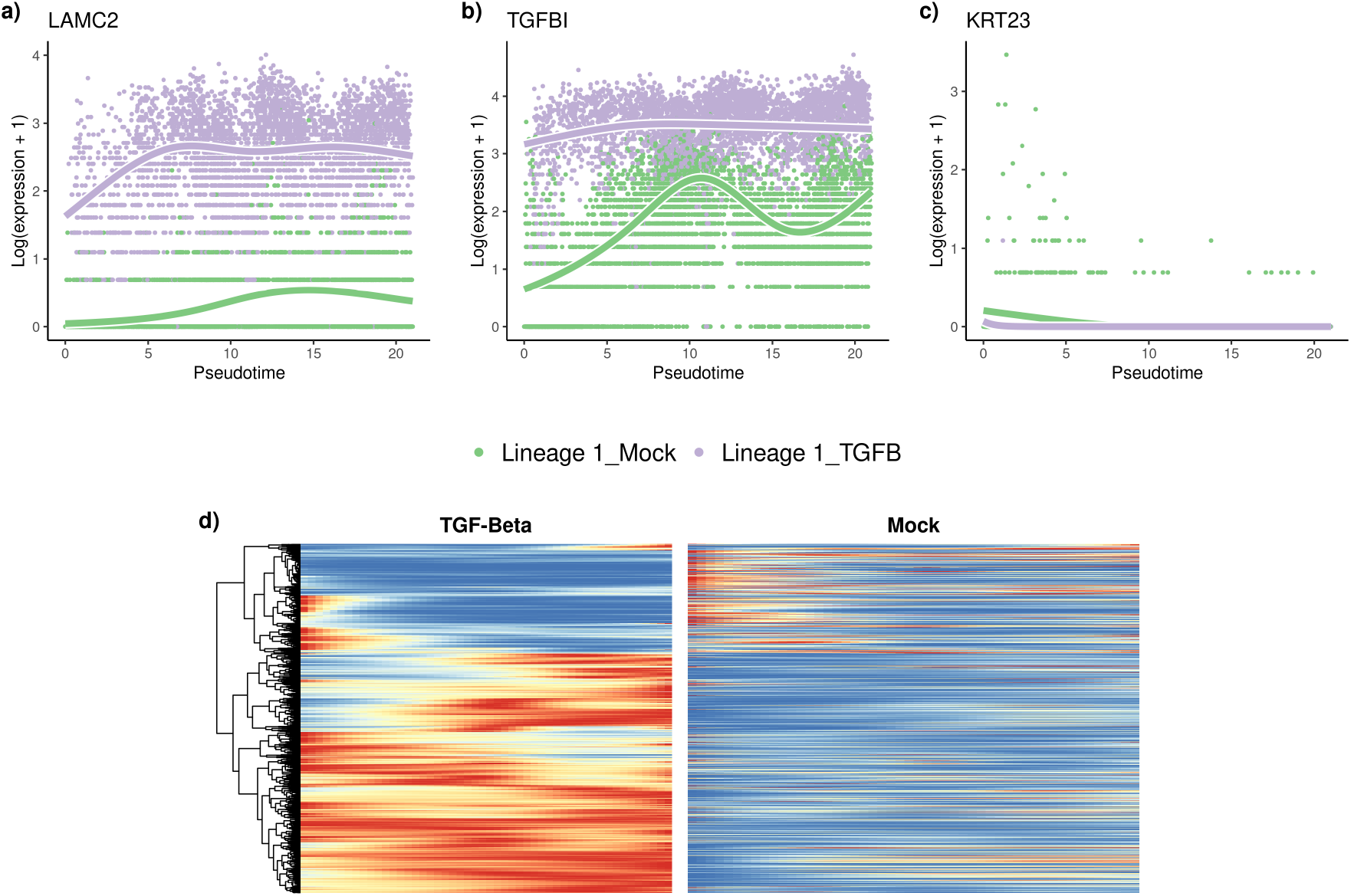
**TGFB** dataset: Differential expression. The `tradeSeq` gene expression model is fitted using the trajectory inferred by `slingshot`. Differential expression between conditions is assessed using the `conditionTest` and genes are ranked according to the test statistics. The genes with the highest **(a.)**, second highest **(b.)**, and smallest **(c.)** test statistics are displayed. After adjusting the *p*-values to control the FDR at a nominal level of 5%, we display genes for both conditions using a pseudocolor image **(d.)** after scaling each gene to a [0,1] range.

Finally, we perform a gene set enrichment analysis on all the genes that are differentially expressed between the conditions. The full results are available in Supplementary Table S1. Top annotations include gene sets involved in cell motility, adhesion, and morphogenesis, which are consistent with the expected biology.

### TCDD dataset

Nault et al. [10] collected a dataset of 16,015 single nuclei to assess the hepatic effects of 2,3,7,8-tetrachlorodibenzo-p-dioxin or TCDD. In particular, they focused on the effect of TCDD on the 9, 951 hepatocytes cells along the central-portal axis. This dataset is not a developmental dataset per se but still exhibits continuous changes along a spatial axis, demonstrating the versatility of the trajectory inference framework in general, and of the `condiments` workflow in particular.

Fig. S3a shows a reduced-dimensional representation of the dataset, with cells labelled according to treatment/control condition, while Fig. S3b shows the same plot colored by cell type, as derived by the authors of the original publication. The cells are aligned in a continuum, from central to mid-central and then mid-portal and portal. The imbalance score shows some spatial pattern (Fig. S3c). However, the nominal p-value associated with the `topologyTest` is .07. We therefore fail to reject the null and we infer a common trajectory using `slingshot` on the spatial clusters. This results in a single-lineage trajectory that respects the ordering of the spatial clusters (Fig. S3d). Note that, since the trajectory reflects a spatial continuum rather than a temporal one, the start of the trajectory is arbitrary. However, inverting the start and end clusters amounts to an affine transformation of the pseudotimes for all the cells. Step 2 and 3 are fully invariant to this transformation, so we can pick the Central cluster as the start of the trajectory.

The densities of the treatment and control pseudotime distributions differ greatly visually (Fig. S3e), with the TCDD density heavily skewed toward the start of the trajectory. Indeed, the `progressionTest` has a nominal p-value ≤ 2.2 × 10^-16^. This coincides with the finding of the original publication which highlighted the periportal hepatotoxicity of TCDD.

The ability of the `progressionTest` to correctly find large-scale changes in the spatial distribution of cells between conditions underscores why we favor fitting a common trajectory. Indeed, the *p*-value of the `topologyTest` in Step 1 is rather small and would have been below .05 if we had not conducted a test against a threshold. However, testing against a threshold and thus fitting a common trajectory does not stop the workflow from finding large-scale differences between conditions in Step 2 and results in a more stable estimate of the trajectory.

After gene filtering, we test 8, 027 genes for spatial differential expression between conditions and we find 2, 114 DE genes when controlling the FDR at a nominal level of 5%. The genes with the largest, second largest, and smallest test statistics are displayed in Fig. S4a-c. Similarly to Nault et al. [10], we obtain a list of zonal genes from Halpern et al. [26]. The proportion of zonal genes among the DE genes is twice their proportion among non-DE genes.

### KRAS datasets

Xue et al. [23] studied the impact of KRAS(G12C) inhibitors at the single-cell level on three models of KRAS(G12C) lung cancers. Specifically, they examined how various cell populations react to these inhibitors and how some cells can return in proliferation mode shortly after the end of the treatment. Here, we want to investigate how the three cancer models (H358, H2122, and SW1573) differ in their response to the KRAS(G12C) inhibitors.

We use the reduced-dimensional representation from the original paper to display the 10, 177 cells from the various types (Fig 5a). Using the cancer type labels and the reduced-dimensional coordinates, an imbalance score can be computed (Fig 5b); some regions clearly show an imbalance. This is further confirmed by the `topologyTest`, with *p*-value smaller than 2.2 × 10^-16^. We therefore do not fit a common trajectory to all cancer types (Fig 5c).

**Figure 5:**
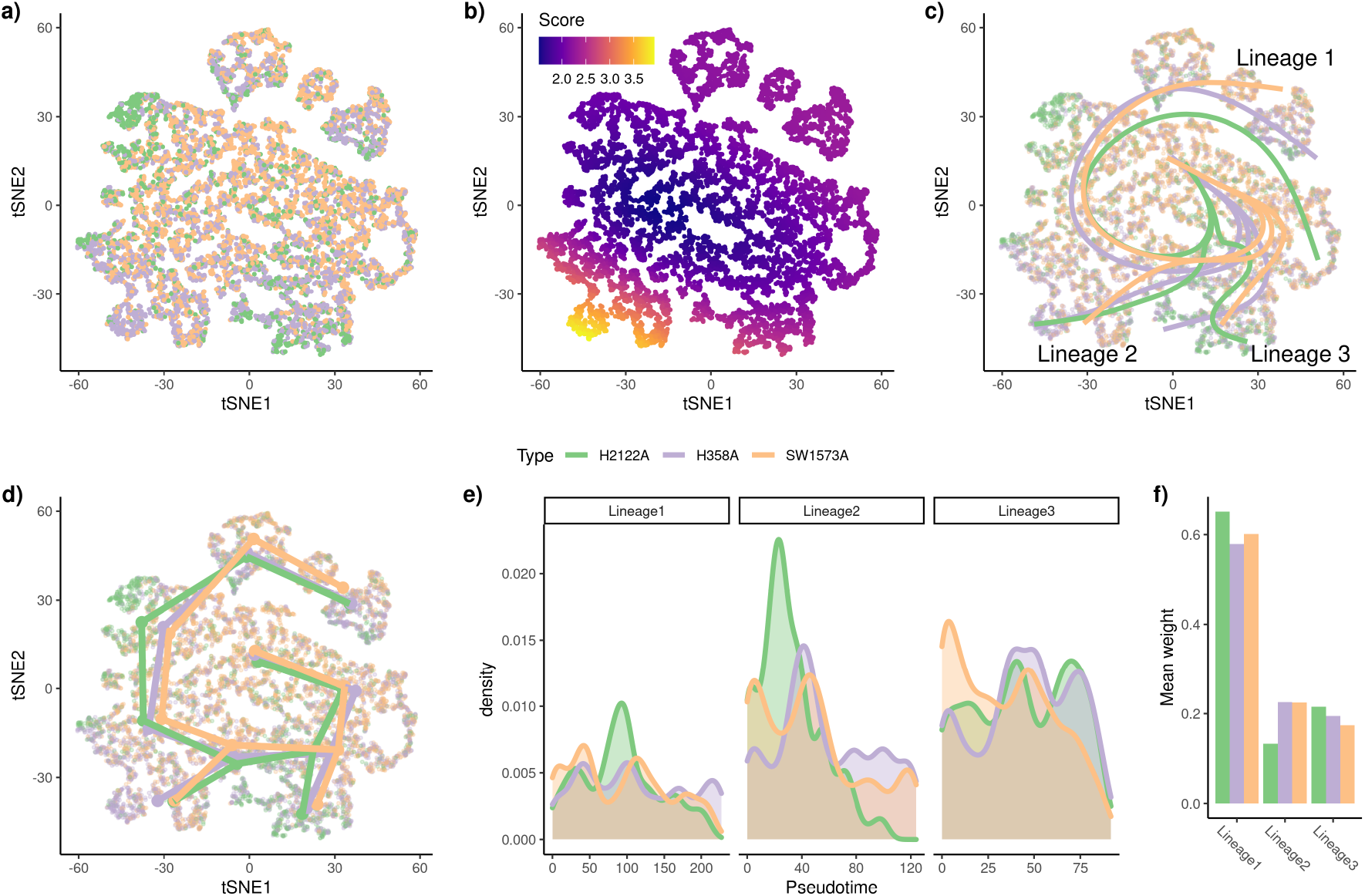
**KRAS** dataset: Differential topology, differential progression, and differential differentiation. Using the reduced-dimensional representation of the original publication (t-SNE), the cells can be colored by cancer type **(a.)**. Using the cancer type label and the reduced-dimensional coordinates, an imbalance score is computed and displayed **(b.)**. The `topologyTest` rejects the null hypothesis of a common trajectory, we thus fit one trajectory per condition **(c.)** . However, the skeleton graphs have the same structure **(d.)**, so we can progress to the next steps in the `condiments` workflow. There is differential progression **(d.)** and we indeed reject the null of identical pseudotime distributions along the trajectory using the `progressionTest`. Similarly, there is differential differentiation **(e.)** and we reject the null of identical weight distributions along the trajectory using the `differentiationTest`. Here, we summarize the distributions by looking at the average weight for each lineage in each condition, which already shows some clear differences.

Note that this does not necessarily imply that the trajectory of reaction to the KRAS(G12C) inhibitors is different between cancer types. Indeed, this may also reflect strong batch effects between conditions, which the normalization scheme was unable to fully remove when integrating the three cancer types in one common reduced-dimensional representation. Thus, it is not really possible to draw a biological conclusion at this first step. However, this does mean that a separate trajectory should be fitted to each condition.

Here, the trajectories, although different, are similar enough that we can still use an underlying common skeleton (Fig 5d). Indeed, we keep the tree structure derived by computing the minimum spanning tree (MST) on the clusters using all cells. This way, it is possible to derive a one-to-one mapping between the lineages of the three trajectories and we respect the assumptions detailed in Section 1.2 that are necessary for the `progressionTest` and `differentiationTest`.

Using this common mapping, we can then proceed to the `progressionTest`. At the global trajectory level, the nominalp-value is smaller than 2.2 × 10^-16^, showing clear differential progression. At the lineage level, all three lineages show strong differential progression, with *p*-values of 2.2 × 10^-16^, 1.2 × 10^-12^, and 1.2 × 10^-14^, respectively. The density plots for the pseudotime distributions at the single-lineage level (Fig 5e) indicate that the differential progression is driven by a group of cells from cancer type H2122A. This matches the top left part of the reduced-dimensional plot, the region where cells exit the initial inhibition stage to enter the reactivation stage. The second lineage also shows a difference between H2122A and the two other models. The pseudotime distribution is heavily skewed toward earlier points in that model compared to the other two. Lineage 2 represents differential progression to a drug-induced state. In Lineage 3, it is the SW1573A model that displays more differential progression.

The `differentiationTest` also has a p-value smaller than 2.2 × 10^-16^. Although all pairwise comparisons are significant, the test statistics are much higher for the Lineage 2 vs. 1 and Lineage 2 vs. 3 comparisons. This again suggests that one model differentiates less into the drug-induced path, compared to the other two. Since the weights have to sum to 1, the 3-dimensional distribution can be fully summarized by any two components. Fig S5 shows clear differences in distributions but visually interpreting different 2D distributions is still challenging. A simpler way to compare the distributions is to look at the average weight in each condition for each lineage (Fig 5f). This ignores the correlation between lineages but still allows for some interpretation. We can see in particular that Lineages 1 and 3 have greater weights for H2122A than for the other two conditions, which is consistent with the different pairwise statistics.

With the mapped trajectories, we can also perform gene-level analysis using the `conditionTest`. When comparing genes across all lineages and conditions, we find 363 differentially expressed genes when controlling the FDR at nominal level 5%. We show the genes with the highest, second highest, and smallest test statistics in Fig. S6a-c. Displaying these global patterns across all three lineages and all three conditions makes it hard to interpret. We therefore focus on the first (and longest) lineage. In that lineage, we find 366 DE genes and we show their expression patterns along Lineage 1 in all three cancer models in Fig. S6d.

## Discussion

In this manuscript, we have introduced `condiments`, a full workflow to analyze dynamic systems under multiple conditions. By separating the analysis into several steps, `condiments` offers a flexible framework with increased interpretability. Indeed, we follow a natural progression through a top-down approach, by first studying overall differences in trajectories with the `topologyTest`, then differences in abundances at the trajectory level with the `progressionTest` and `differentiationTest`, and finally gene-level differences in expression with the conditionTest.

As demonstrated in the simulation studies, taking into account the dynamic nature of systems via the trajectory representation enables `condiments` to better detect true changes between conditions. The flexibility offered by our implementation, which provides multiple tests for non-parametric comparisons of distributions, also allows us to investigate a wide array of scenarios. This is evident in the three case studies presented in the manuscript. Indeed, in the first case study we have a developmental system under treatment and control conditions, while in the second case study the continuum does not represent a developmental process but spatial separation. In the third case study, the conditions do not reflect different treatments but instead different cancer models. This shows that `condiments` can be used to analyze a wide range of datasets.

Often, the different conditions also represent different batches. Indeed, some interventions cannot be delivered on a cell-by-cell basis and this creates unavoidable confounding between batches and conditions. Normalization and integration of the datasets must therefore be done without eliminating the underlying biological signal. This balance can be hard to strike, as discussed in Zhao et al. [17]. Proper experimental design – such as having several batches per condition – or limiting batch effects as much as possible – for example, sequencing a mix of conditions together – can help lessen this impact. Still, some amount of confounding is sometimes inherent to the nature of the problem under study.

The tests used in the workflow (e.g., Kolmogorov-Smirnov test) assume that the pseudotime and weight vector are known and independent observations for each cell. However, this is not the case: they are estimated using TI methods which use all samples to infer the trajectory, and each estimate inherently has some uncertainty. Here, we ignore this dependence, as is the case in other differential abundance methods, which assume that the reduced-dimensional coordinates are observed independent random variables even thought they are being estimated using the full dataset. We stress that, rather than attaching strong probabilistic interpretations to p-values (which, as in most RNA-seq applications, would involve a variety of hard-to-verify assumptions and would not necessarily add much value to the analysis), we view the p-values produced by the `condiments` workflow as useful numerical summaries for guiding the decision to fit a common trajectory or condition-specific trajectories and for exploring trajectories across conditions and identifying genes for further inspection.

Splitting the data into two groups, where the first is used to estimate the trajectory and the second is used for pseudotime and weight estimation could, in theory, alleviate the dependence issue, at the cost of smaller sample sizes. However, this would ignore the fact that, in practice, users perform exploratory steps using the full data before performing the final integration, dimensionality reduction, and trajectory inference. Moreover, results on simulations show that all methods considered keep excellent control of the false discovery rate despite the violation of the independence assumptions. This issue of “double-dipping” therefore seems to have a limited impact.

The two issues raised in the previous paragraphs highlight the need for independent benchmarking. Simulation frameworks such as `dyngen` [22] are crucial. They also need to be complemented by real-world case studies, which will become easier as more and more datasets that study dynamic systems under multiple conditions are being published. `condiments` has thus been developed to be a general and flexible workflow that will be of use to researchers asking complex and ever-changing questions.

## Acknowledgments

KVdB is a postdoctoral fellow of the Belgian American Educational Foundation (BAEF) and is supported by the Research Foundation Flanders (FWO), grants 1246220N and G062219N.

## Authors’ contributions

### Data and code availability

The results from this paper can be fully reproduced by following along the vignettes at https://hectorrdb.github.io/condimentsPaper. These also contain the code needed to recreate the datasets used for the simulations, as well as processed versions of all three datasets used in the case studies, augmented by metadata and functions to recreate the processed versions, using raw counts obtained from GEO (**TGFB** dataset: GSE114687, **TCDD** dataset: GSE148339, **KRAS** dataset: GSE137912).

The `condiments` workflow is available as an R package from Github (https://github.com/HectorRDB/condiments) and will be made available through the Bioconductor Project.

All the methods to test for equality of two (or *k*) distributions have been put together for use by others in an R package called `Ecume`, available through CRAN and that can be explored at https://hectorrdb.github.io/Ecume.

## Methods

### 1.1 Tests for equality of distributions

#### 1.1.1 General setting

Consider a set of n i.i.d. observations, **X**, with **X**_*i*_ ~ **P**_1_, and a second set of m i.i.d. observations, **Y**, with **Y**_*j*_ ~ **P**_2_, independent from **X**. For example, in our setting, **X** and **Y** may represent estimated pseudotimes for cells from two different conditions. We limit ourselves to the case where **X** and **Y** are random vectors of the same dimension *d*.

The general goal is to test the null hypothesis that **X** and **Y** have the same distribution, i.e., *H*_0_: **P**_1_ = **P**_2_.

#### 1.1.2 Univariate case: The weighted Kolmogorov-Smirnov test

##### The two-sample Kolmogorov-Smirnov test

Consider the case where **X**_*i*_ and **Y***j* are scalar random variables (i.e., *d* = 1). The associated empirical cumulative distribution functions (ECDFs) are denoted, respectively, by **F**_1,*n*_ and **F**_2,*m*_. The univariate case occurs, for example, when there is only one lineage in the trajectory(ies), so that the pseudotime estimates are scalars.

In this setting, one can test *H*_0_ using the standard Kolmogorov-Smirnov test [19], with test statistic defined as:

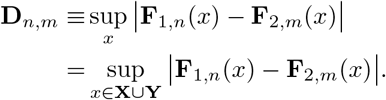

The rejection region at nominal level *α* is

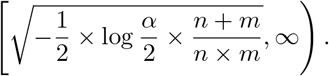

That is, we reject the null hypothesis at the *α*-level if and only if 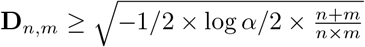.

##### The two-sample weighted Kolmogorov-Smirnov test

Consider a more general setting where we have weights *w*_1,*i*_ ∈ [0,1] and *w*_2,*j*_ ∈ [0,1] for each of the observations. In trajectory inference, the weights may denote the probability that a cell belongs to a particular lineage in the trajectory. Following Monahan [27], we modify the Kolmogorov-Smirnov test in two ways. Firstly, the empirical cumulative distribution functions are modified to account for the weights

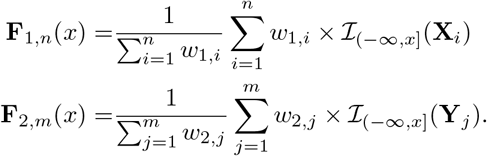

Secondly, the definition of **D**_*n,m*_ is unchanged, but the significance threshold is updated, that is, the rejection region is

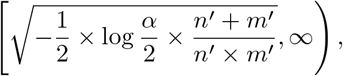

where

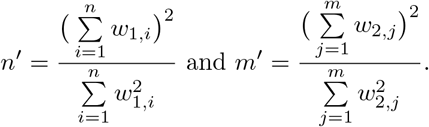

#### 1.1.3 Multivariate case: The classifier test

##### Concept

Suppose that we have a classifier *δ*(·), which could be, for example, a multinomial regression or SVM classifier. This classifier is a function from the support of **X** and **Y** into {1, 2}. The data are first split into a learning and a test set, such that the test set contains *n*_test_ observations, equally-drawn from each population, i.e., there are *n*_test_/2 observations **X**^(test)^ from **X** and *n*_test_/2 observations **Y**^(test)^ from **Y**. Next, the classifier is trained on the learning set. We denote by 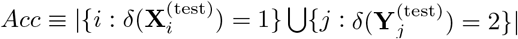 the number of correct assignations made by the classifier on the test set.

If *n = m*, under the null hypothesis of identical distributions, no classifier will be able to perform better on the test set than a random assignment would, i.e., where the predicted label is a *Bernoulli(1/2)* random variable. Therefore, testing the equality of the distributions of **X** and **Y** can be formulated as testing

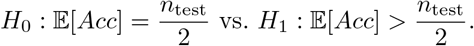

Under the null hypothesis, the distribution of Acc is:

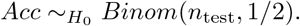

As detailed in Lopez-Paz and Oquab [20], one can use the classifier to devise a test that will guarantee the control of the Type 1 error rate.

##### The classifier test in practice

In practice, we make no assumptions about the way in which the distributions we want to compare might differ, which means the classifier needs to be quite flexible. Following Lopez-Paz and Oquab [20], we chose to use either a k-nearest neighbor classifier (*k*-NN) or a random forests classifier [28], since such classifiers are fast and flexible. Hyper-paramters are chosen through cross-validation on the learning set. To avoid issues with class imbalance, we downsample the distribution with the largest number of samples first so that each distribution has the same number of observations. That is, we have *n*’ = min(*m,n*) observations in each condition (or class). A fraction (by default 30%, user-defined) is kept as test data, so that *n*_test_ = .3 × *n*’. We then train the classifier on the learning data, and select the tuning parameters through cross-validation on that learning set. Finally, we predict the labels on the test set and compute the accuracy of the classifier on that test set. This yields our classifier test statistic.

##### Power of the classifier test

It is interesting to note that the classifier test is valid no matter the classifier chosen. However, the choice of classifier will have obvious impact on the power of the test.

#### 1.1.4 Multivariate case: Other methods

Although we have found that the classifier test performs best in practice, there are many methods that test for the equality of two multivariate distributions. We have implemented a few such methods in `condiments`, in case users would like to try them: The two-sample kernel test [29] and the permutation test relying on the Wasserstein distance (see descriptions in the supplementary methods).

#### 1.1.5 Extending the setting by considering more than two conditions

Consider *C* ≥ 2 sets of samples, such that, for *c* ∈ {1,…, *C*}, we have *n_c_* i.i.d. observations **X**^(*c*)^ with 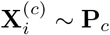. We want to test the null hypothesis:

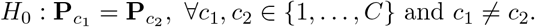

While extensions of the Kolmogorov-Smirnov test [30] and the two-sample kernel test [31] have been proposed, we choose to focus only on the framework that is most easily extended to *C* conditions, namely, the classifier test. Indeed, the *C*-condition classifier test requires choosing a multiple-class classifier instead of a binary classifier (which is the case for the *k*-NN classifier and random forests), selecting *n*_test_/*C* observations for each class in the test set, and testing:

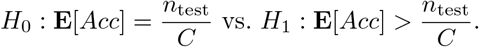

Under the null distributions, the distribution of *Acc* is:

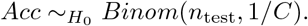

#### 1.1.6 Extending the setting by considering an effect size

##### Effect size for the Kolmogorov-Smirnov test

The null hypothesis of the (weighted) Kolmogorov-Smirnov test is *H*_o_: *P*_1_ = *P*_2_. We can modify this null hypothesis by considering an effect size threshold *t*, such that *H*_0_(*t*): sup_*x*_ |*P*_1_(*x*) — *P*_2_(*x*)| ≤ *t*. The test statistic is then modified as:

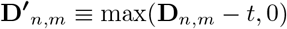

and the remainder of the testing procedure is left unchanged.

##### Effect size for the classifier test

Similarly, the null and alternative hypotheses of the classifier test can be modified to test against an effect size threshold *t* as follows

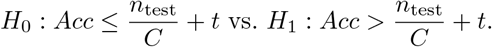

### 1.2 General statistical model for the trajectories

Consider a set of condition labels *c* ∈ {1, …,*C*} (e.g.,”treatment” or “control”, “knock-out” or “wildtype”). For each condition, there is a given topology/trajectory 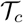 that underlies the developmental process. This topology is generally in the form of a tree, with a starting state which then differentiates along one or more lineages; but one can also have a circular graph, e.g., for the cell cycle. In general, a trajectory is defined as a directed graph.

We denote by *L_c_* the number of unique paths – or lineages – in the trajectory 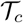 and by 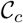 the set of cells that belong to condition c. For example, for a tree structure, paths go from the root node (stem cell type) to the leaf nodes (differentiated cell type). For a cell cycle, any node can be be used as the start. A cell *i* from condition *c_i_* is characterized by the following features:

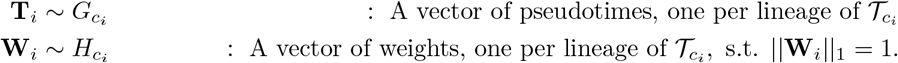

Note that the distribution functions are condition-specific. We further make the following assumptions:

- All *G_c_* and *H_c_* distributions are continuous;
- The support of all *G_c_* is bounded in ℝ*^L_c_^*;
- The support of all *H_c_* is [0,1]*^L_c_^*.

The gene expression model will be discussed below, in the differential expression section.

#### Trajectory inference

Many algorithms have been developed to estimate lineages from single-cell data [5]. Most algorithms provide a binary indicator of lineage assignment, that is, the **W**_*i*_ vectors are composed of 0s and 1s, so that a cell either belongs to a lineage or it does not (note that when cells fall along a lineage prior to a branching event, this vector may include multiple 1s, violating our constraint that the **W**_*i*_ have unit norm. In such cases, we normalize the weights to sum to 1).

#### Mapping between trajectories

Many of the tests that we introduce below assume that the cells from different conditions follow “similar” trajectories. In practice, this means that we either have a common trajectory for all conditions or that there is a possible manual mapping from one lineage to another. The term “mapping” is more rigorously defined as follows.

##### Definition 1

The trajectories 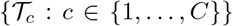 have a mapping if and only if ∀(c_1_,c_2_) ∈ {1,…,C}^2^, 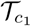 and 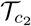 are isomorphic.

If there is a mapping, this implies in particular that the number of lineages L_c_ per trajectory *T_c_* is the same across all conditions c and we call this this value L. Since a graph is always isomorphic with itself, a common trajectory is a special case of a situation where there is a mapping.

##### Definition 2

The trajectories 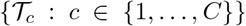 have a partial mapping if and only if ∀(c_1_,c_2_) ∈ {1,…,C}^2^, there is a subgraph 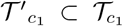 and a subgraph 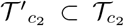 that are isomorphic.

Essentially, this means that the size of the changes induced by the various conditions do not disturb the topology of the original trajectory *too much*. The above mathematical definitions aim to formalize what *too much* is. Indeed, if the conditions lead to very drastic changes, it will be quite obvious that the trajectories are different and comparing them will mostly be either non-informative or will not require a complex framework. We aim to build a test that retains reasonable power in more subtle cases.

### 1.3 Differential topology

#### Imbalance score

Consider a set of n cells, with associated condition labels *c_i_* ∈ {1, …,*C*} and coordinate vectors **X**_*i*_ in *d* dimensions, usually corresponding to a reduced-dimensional representation of the expression data obtained via PCA or UMAP[24, 32].

Let **p** = {*p_c_*}_*c*∈{1,…,*c*}_ denote the “global” distribution of cell conditions, where *p_c_* is the overall proportion of cells with label *c* in the sample of size *n*. The imbalance score of a cell reflects the deviation of the “local” distribution of conditions in a neighborhood of the cell compared to the global distribution **p**. Specifically, for each cell *i*, we compute its *k*-nearest neighbor graph using the Euclidean distance in the reduced-dimensional space. We therefore have a set of *k* neighbors and a set of associated neighbor condition labels *c_i,κ_* for *κ* ∈ {1,…, *k*}. We then assign to the cell a *z*-score, based on the multinomial test statistic 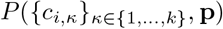, as defined in Section S-1.2. Finally, we smooth the z-scores in the reduced-dimensional space by fitting s cubic splines for each dimension. The fitted values for each of the cells are the imbalance scores. Thus, the imbalance scores rely on two user-defined parameters, *k* and s. We set default values of 10 for both parameters. However, since this is meant to be an exploratory tool, we encourage users try different values for these parameters and observe the changes to better understand their data.

#### General setting for the topologyTest

The imbalance score only provides a qualitative visual inspection of local imbalances in the distribution of cell conditions. However, we need a more global and formal way to test for differences in topology between condition-specific trajectories. That is, we wish to test the null hypothesis

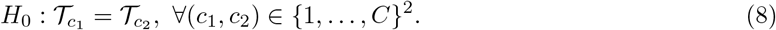

In practice, in order to test *H*_0_, we have a set of cells *i* with condition labels *c_i_*. We can estimates the pseudotimes of each cell when fitting a trajectory for each condition. We then want to compare this distribution of pseudotimes to a null distribution. To generate this null distribution, we use permutations in the following manner

a. Estimate **T**_*i*_ for all *i* by inferring one trajectory per condition, using any trajectory inference method.
b. Randomly permute the condition labels *c_i_* to obtain new labels 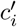, re-estimate **T**’_*i*_ for each *i*.
c. Repeat the permutation *r* times (by default, *r* = 100).

Under the null hypothesis, the *n* **T**_*i*_ should therefore be drawn from the same distribution as the *r × n* **T**’_*i*_. We can test this using the weighted Kolmogorov-Smirnov test (if *L* = 1), the kernel two-sample test (if *C* = 2), or the classifier test (any C). This is the `topologyTest`.

The aforementioned tests require that the samples be independent between the two distributions under comparison. However, here, the two distributions correspond to different pseudotime estimates for the same cells so the samples are not independent between distributions. Even within distributions, the independence assumption is violated: the pseudotimes are estimated using trajectory inference methods that rely on all samples. Moreover, within the **T**’_*i*_, we have *r* pseudotime estimates of each cell.

The first two violations of the assumptions are hard to avoid and are further addressed in the discussion section. However, we can eliminate the third one by simply taking the average **T**’_*i*_ for each cell. We then compare two distributions each with n samples. Both options (with and without averaging) are implemented in the `condiments` R package, but the default is the average.

Furthermore, rather than attaching strong probabilistic interpretations to *p*-values (which, as in most RNA-seq applications, would involve a variety of hard-to-verify assumptions and would not necessarily add much value to the analysis), we view the p-values produced by the `condiments` workflow simply as useful numerical summaries for exploring trajectories across conditions and identifying genes for further inspection.

#### Running the topologyTest in practice

Under the null, there should exist a mapping between trajectories, both within conditions and between permutations. However, in practice, most trajectory inference methods will be too unstable to allow for automatic mapping between the runs. Indeed, they might find a different number of lineages for some runs. Moreover, even if the number of lineages and graph structure remained the same across all permutations, mapping between permutations would break even more the independence assumption since the condition labels would need to be used.

Therefore, for now, the `topologyTest` test is limited to two trajectory inference methods, slingshot [2] and TSCAN [4], where a set graph structure can be prespecified. Both methods rely on constructing a minimum spanning tree (MST) on the centroids of input clusters in a reduced-dimensional space to model branching lineages. In TSCAN, a cell’s pseudotime along a lineage is determined by its projection onto a particular branch of the tree, and its weight of assignment is determined by its distance from the branch. `slingshot` additionally fits simultaneous principal curves. A cell’s pseudotime along a lineage is determined by its projection onto a particular curve and its weight of assignment is determined by its distance from the curve. We therefore construct the MST on the full dataset (i.e., using all the cells regardless of their condition label), based on user-defined cluster labels. Then, we keep the same graph structure as input to either TI method: the nodes are the centers of the clusters, but restrained to cells of a given condition. This way, the path and graph structure are preserved. Note however, that there no guarantee that the graph remains the MST when it is used for TI on a subset of cells.

### 1.4 Testing for differential progression

The differential progression test requires that a (partial) mapping exists between trajectories. If the mapping is only partial, we restrict ourselves to the mappable parts of the trajectories (i.e., subgraphs).

#### Testing for differential progression for a single lineage

For a given lineage *l*, we want to test the null hypothesis that the pseudotimes along the lineage are identically distributed between conditions, which we call *identical progression.* Following the above notation, we want to test that the *l^th^* components *G_lc_* of the distribution functions *G_c_* are identical across conditions

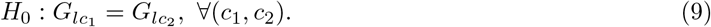

#### Testing for global differential progression

We can also test for global differences across all lineages, that is,

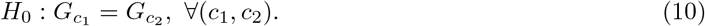

#### Possible tests

If *C* = 2, all tests introduced in Section 1.1 can be used to test the hypothesis in Equation (9). If *C* > 2, we need to rely on the classifier test.

If *L* = 1, the hypotheses in Equations (9) and (10) are identical. However, for *L* > 1, the functions *G_c_* are not univariate distributions.

#### Using the Kolmogorov-Smirnov test in the *L* > 1 setting

For *L* > 1, we can use lineage-level weights as observational weights for each individual lineage, which is an appealing property. Two settings are possible.

- Test the null hypothesis in Equation (9) for each lineage using the Kolmogorov-Smirnov test and perform a global test using the classifier test or the kernel two-sample test.
- Test the null hypothesis in Equation (9) for each lineages using the Kolmogorov-Smirnov test and combine the *p*-values *p_l_* for each lineage *l* using Stouffer’s Z-score method [33], where each lineage is associated with observational weights 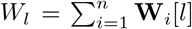. The nominal *p*-value associated with the global test is then

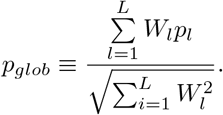

Note that the second setting violates the assumption of Stouffer’s Z-score method, since the *p*-values are not i.i.d. However, this violation does not seem to matter in practice and this test outperforms others so we set it as default.

### 1.5 Testing for differential differentiation

The differential progression test requires that a (partial) mapping exists between trajectories. If the mapping is only partial, we restrict ourselves to the mappable parts of the trajectories.

#### Testing for differential differentiation for a single pair of lineages

For a given pair of lineages *l, l*’, we want to test the null hypothesis that the cells differentiate between *l* and *l*’ in the same way between all conditions, which we call *identical differentiation.* Following the above notation, we want to test that the *l^th^* and *l’^th^* components of the distibution function *H_c_* are the same

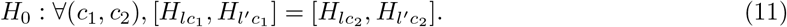

#### Testing for global differential differentiation

We can also test for a global difference across all pairs of lineages, that is,

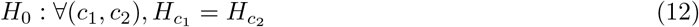

#### Possible tests

Since all variables are multivariate, we cannot use the Kolmogorov-Smirnov test. By default, this test relies on the classifier test with random forest as a classifier.

### 1.6 Testing for differential expression

#### Notation

The gene expression model does not require a mapping or even a partial mapping. Indeed, it can work as well with a common trajectory, different trajectories, or even a mix where some lineages can be mapped between the trajectories for various conditions and others cannot. To reflect this, we consider all *L_tot_* lineages together. We introduce a new weight for each cell

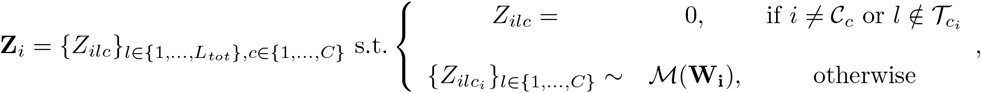

where 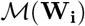 is a binary (or one-hot) encoding representation of a multinomial distribution with proportions **W**_*i*_ as in `tradeSeq`.

Likewise, we modify the pseudotime vector to have length *L_tot_* such that

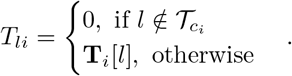

#### Gene expression model

We adapt the model from Van den Berge et al. [8] to allow for condition-specific expression. For a given gene *j*, the expression measure *Y_ji_* for that gene in cell *i* can be modeled thus:

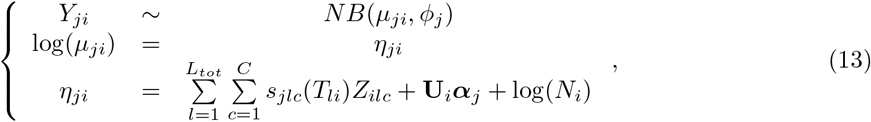

where the mean *μ_ji_* of the negative binomial distribution is linked to the additive predictor *η_ji_* using a logarithmic link function. The **U** matrix represents an additional design matrix, representing, for example, a batch effect.

The model relies on lineage-specific, gene-specific, and condition-specific smoothers *s_jlc_*, which are linear combinations of *K* cubic basis functions, 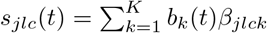.

#### Testing for differential Expression

With this notation, we can introduce the `conditionTest`, which, for a given gene *j*, tests the null hypothesis that the smoothers are identical across conditions:

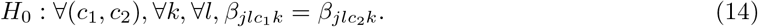

We fit the model using the `mgcv` package [34] and test the null hypothesis using a Wald test for each gene. Note that, although the gene expression model can be fitted without any mapping, the `conditionTest` only exists for lineages with at least a mapping for two conditions.

### 1.7 Simulation study

#### 1.7.1 Simulating datasets

The simulation study relies on the dyngen framework of Cannoodt et al. [22] and all datasets are simulated as follows. 1/ *A* common trajectory is generated, with an underlying gene network that drives the differentiation along the trajectory. 2/ *A* set of *N_WT_* cells belonging to the wild-type condition (i.e., with no modification of the gene network) is generated. 3/ One master regulator that drives differentiation into one of the lineages is impacted, by multiplying the wild-type expression rate of that gene by a factor *m*. If *m* =1, there is no effect; if *m* > 1, the gene is over-expressed; and if *m* < 1, the gene is under-expressed, with *m* = 0 amounting to a total knock-out. 4/ *A* set of *N_KO_* = *N_WT_* cells is generated using the common trajectory with the modified gene network. 5/ *A* common reduced-dimensional representation is computed.

We generate three types of datasets, over a range of values of *m*: a simple trajectory with *L* = 2 lineages and *C* = 2 conditions (WT and KO) named 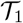; a trajectory with two consecutive branchings with *L* = 3 lineages and *C* = 2 conditions (WT and KO) named 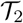; and a simple trajectory with *L* = 2 lineages and *C* = 3 conditions (WT, KO, and UP) named 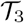. For the latter case, Steps 3-4/ are repeated twice, with values of *m* for KO and 1/*m* for UP.

For 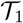 and 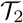, we use values of *m* ∈ {.5, .8, .9, .95,1,1/.95,1/.9,1/.8,1/.5}, such that at the extremes the KO cells fully ignore some lineages. Values of .95 and 1/.95 represent the closest to no condition effect (*m* = 1), where the effect was still picked out by some tests. For 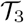, since the simulation is symmetrical in *m*, we pick *m* ∈ {.5, .8, .9, .95,1}. We have one large dataset per value of *m* and per trajectory type. We use those large datasets to generate smaller ones of size *n*, by sampling 10% of the cells from each condition 50 times and applying the various tests on the smaller datasets. The reason for first generating a large dataset and then smaller ones by subsampling instead of generating small ones straightaway are computational: the generation of the datasets is time-consuming and the part that scales with *N_WT_* can be parallelized. Hence, it is almost as fast to generate a large dataset than a small one with `dyngen`. We pick *N_WT_* = 20,000 (for the large dataset) and thus *n* = 2,000.

Since we generate many datasets with true effect (*m* ≠1) but only one null dataset, the size of *N_WT_* for *m* = 1 is doubled to 40, 000. To be comparable, the fraction of cells sampled is decreased to 5% so that *n* = 2, 000 and we perform 100 subsampling. Table 2 recapitulates all this.

**Table 2:** Summary of all simulated datasets. We report the name, number of cells *n_WT_* for values of *m* ≠1 and *m* = 1, number of conditions *C*, number of lineages *L*, impacted master regulator, and figure numbers for the associated gene network and an example of low-dimensional representation.

| Dataset | $N_{WT}$ | | $n$ | $L$ | $C$ | Impacted Regulator | Gene Network Figure | Reduced Dimension Representation |
| --- | --- | --- | --- | --- | --- | --- | --- | --- |
| | $m \neq 1$ | $m = 1$ | | | | | | |
| $\mathcal{T}_1$ | 20,000 | 40,000 | 2,000 | 2 | 2 | B3 | Fig S1a | Fig.2a |
| $\mathcal{T}_2$ | 20,000 | 40,000 | 2,000 | 3 | 2 | D2 | Fig S1b | Fig.2b |
| $\mathcal{T}_3$ | 20,000 | 40,000 | 2,000 | 2 | 3 | B3 | Fig S1a | Fig.2c |

#### 1.7.2 Measuring the performance of the tests on the simulated datasets

To run the `condiments` workflow, we first estimate the trajectories using `slingshot` with the clusters provided by `dyngen`. Then, we run the `progressionTest` and the `differentiationTest` with default arguments.

We compare `condiments` to two other methods. `milo` [16] and `DAseq`[17] both look at differences in proportions within local neighborhoods, using k-nearest neighbor graphs to define this locality. Then, `milo` uses a negative binomial GLM to compare counts for each neighborhood, while `DAseq` uses a logistic classifier test. Therefore, both methods test for differential abundance in multiple regions. To account for multiple testing, we adjust the p-values using the Benjamini, Yoav; Hochberg [21] FDR-controlling procedure.

We select two adjusted p-value cutoffs, .01 and .05, which amount to controlling the FDR at nominal level 1% and 5%, respectively. For a given cutoff *c* and a given dataset, we can look at the results of each test on all simulated datasets for all values of *m*. For each test, the number of true positives (TP) is the number of simulated datasets where *m* ≠ 1 and the adjusted *p*-value is smaller than *c*, the number of true negatives (TN) is the number of simulated datasets where *m* = 1 and the adjusted *p*-value is larger than *c*, the number of false positives (FP) is the number of simulated datasets where *m* =1 and the adjusted *p*-value is smaller than *c*, and the number of false negatives (FN) is the number of subsampled datasets where *m* ≠ 1 and the adjusted *p*-value is larger than *c*. We then examine 5 metrics built on these four variables:

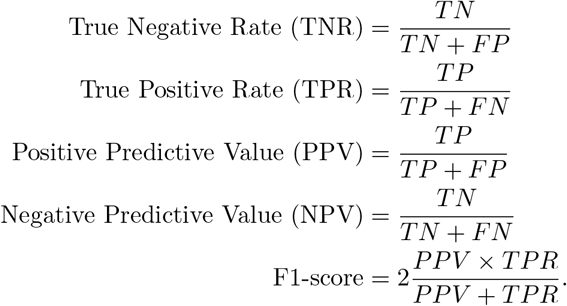

## S-1 Supplementary methods

### S-1.1 Other methods implemented in the condiments package to test equality of distributions

These methods were found to be less efficient in initial benchmarking, but are implemented in case users want to use them.

#### S-1.1.1 Multivariate case: The two-sample kernel test

##### Mean maximum discrepancy

The two-sample kernel test was defined by Gretton et al. [29] and relies on the mean maximum discrepancy (MMD). Considering a kernel function

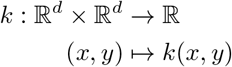

the MMD is then defined as

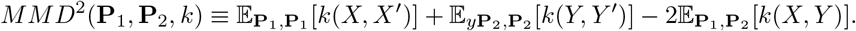

For a properly defined kernel, we have *MMD*^2^ (**P**_1_, **P**_2_,*k*) = 0 i.i.f. **P**_1_ = **P**_2_.

##### Unbiased statistic

Following Gretton et al. [29], we define the unbiased MMD statistic:

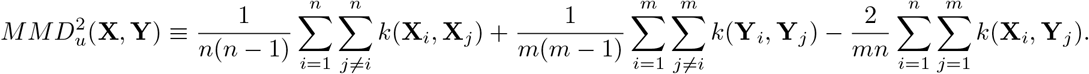

##### Linear statistic for faster computations

While the *MMD*^2^ offers fast convergence, it can be burdensome to compute when *m* and *n* get large. Gretton et al. [29] propose a linear statistic in the case *m = n*. We can extend this in the general setting by just sampling a fixed fraction of the terms of each sum. This lowers kernel computation costs drastically.

##### Null distribution of the statistic

For some kernels, the 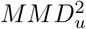 follows some theoretical inequalities under the null that allows one to define rejection regions. However, this is not always the case. Therefore, in practice, we instead rely on permutations to compute a null distribution for the test statistic. Under the null, **X**_*i*_ and **Y**_*j*_ are from the same distribution so they can be swapped in the sums. We can therefore generate an empirical distribution and use it to define rejection regions.

#### S-1.1.2 Multivariate case: Optimal transport

We consider the Wasserstein distance [35, 36], also known as earth’s mover distance, between the two distributions, estimated using the samples **X** and **Y**. We can generate a null distribution for this metric by permuting observations in the combined **X** and **Y** datasets, thereby obtaining a valid test for H_0_: **P**_1_ = **P**_2_. This works in any number of dimensions, but is limited to the two-sample case.

### S-1.2 Mutinomial test

We consider a set of categories arbitrarily numbered from 1 to *C*. Additionally, we consider a null distribution **C**_o_, defined on 1 to *C* by a vector of probabilities 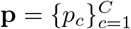. Then, given a set of *n* i.i.d. realizations (*c*(1),…, *c*(*n*)) of a random variable **C**, we can test the null hypothesis *H*_0_: **C** ~ **C**_0_ or, equivalently, *H*_0_: **P**(**C** = *c*) = *p_c_*, ∀_*c*_ ∈ {1,…, *C*}. Under the null, **P**(*c_i_*) = *p_c_i__* and the associated p-value of the multinomial test can be defined as:

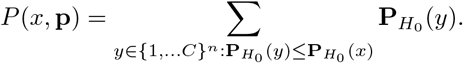

It verifies: ∀*α* ∈ [0: 1], **P**_*H*_0__ (*P*(*x*, **p**) ≤ *α*) ≤ *α*.

### S-1.3 Case studies: Processing

#### TGFB

The two conditions are normalized separately using `SCTransform` [37] and then integrated using `Seurat` [38]. The reduced-dimensional representation is computed using UMAP [24] on the top 50 principal components (PC). The imbalance score is computed with parameters *k* = 20 and *smooth* = 40. The `trajectory` is estimated using slingshot. The `topologyTest` is run with 100 permutations with the Kolmogorov-Smirnov test and default `threshold` of .01. The `progressionTest` is run with defaults. All genes with at least 2 reads in 15 cells are kept. The smoothers are fitted for each gene using 7 knots as recommended by the `evaluateK` function. Gene set enrichment analysis is done using the `fgsea` [39] package on the GO Biological Process ontology sets.

#### TCCD

The dataset is first filtered using the cell type assignments from the original publication to only retains cells labelled as hepatocytes. The count matrix is scaled using `Seurat` [38] and reduced-dimensional coordinates are computed using UMAP [24] on the top 30 PCs. The imbalance score is computed with default *k* and `smooth` = 5. The trajectory is estimated using `slingshot`. The `topologyTest` is run with 100 permutations with the Kolmogorov-Smirnov test and default threshold of .01. The progressionTest is run with defaults. All genes with at least 2 reads in 15 cells are kept; all genes with at least 3 reads in 10 cells are kept. The smoothers are fitted for each gene using 7 knots as recommended by the `evaluateK` function.

#### KRAS

The reduced-dimensional coordinates were obtained from the original publication. The imbalance score is run with defaults and the `topologyTest` is run with 100 permutations with the classifier test and default threshold of .01. The trajectories are estimated using slingshot with parameters *reweight = FALSE* and *reassign = FALSE*. The `progressionTest` and `differentiationTest` are run with defaults. All genes with at least 5 reads in 10 cells are kept. The smoothers are fitted for each gene using 6 knots as recommended by the evaluateK function.

## 1.2 Supplementary figures

### S-2.1 Simulations

**Figure S1:**
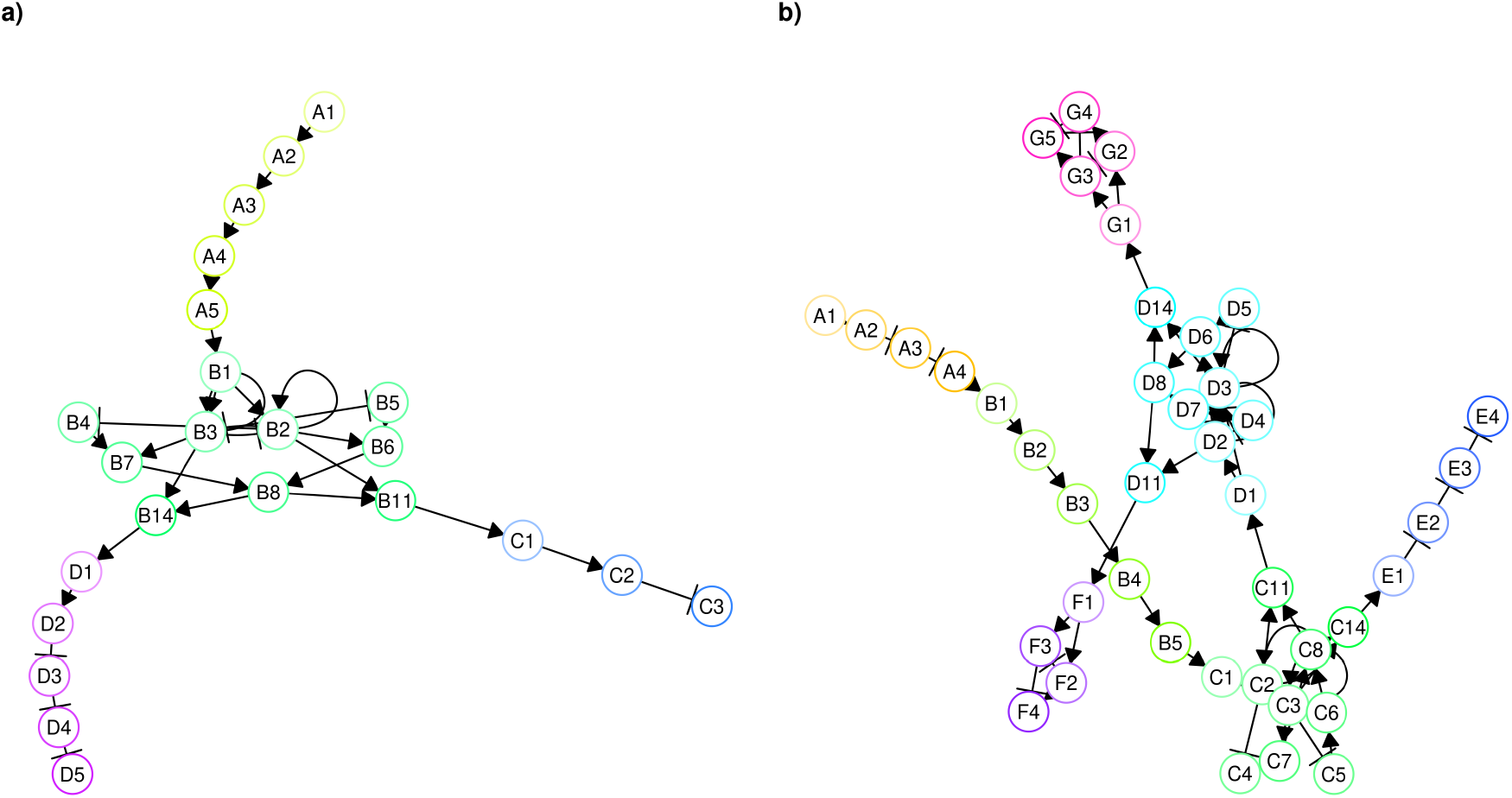
Simulation example. Regulator networks for the **(a.)** two-lineage and **(b.)** three-lineage trajectories.

**Figure S2:**
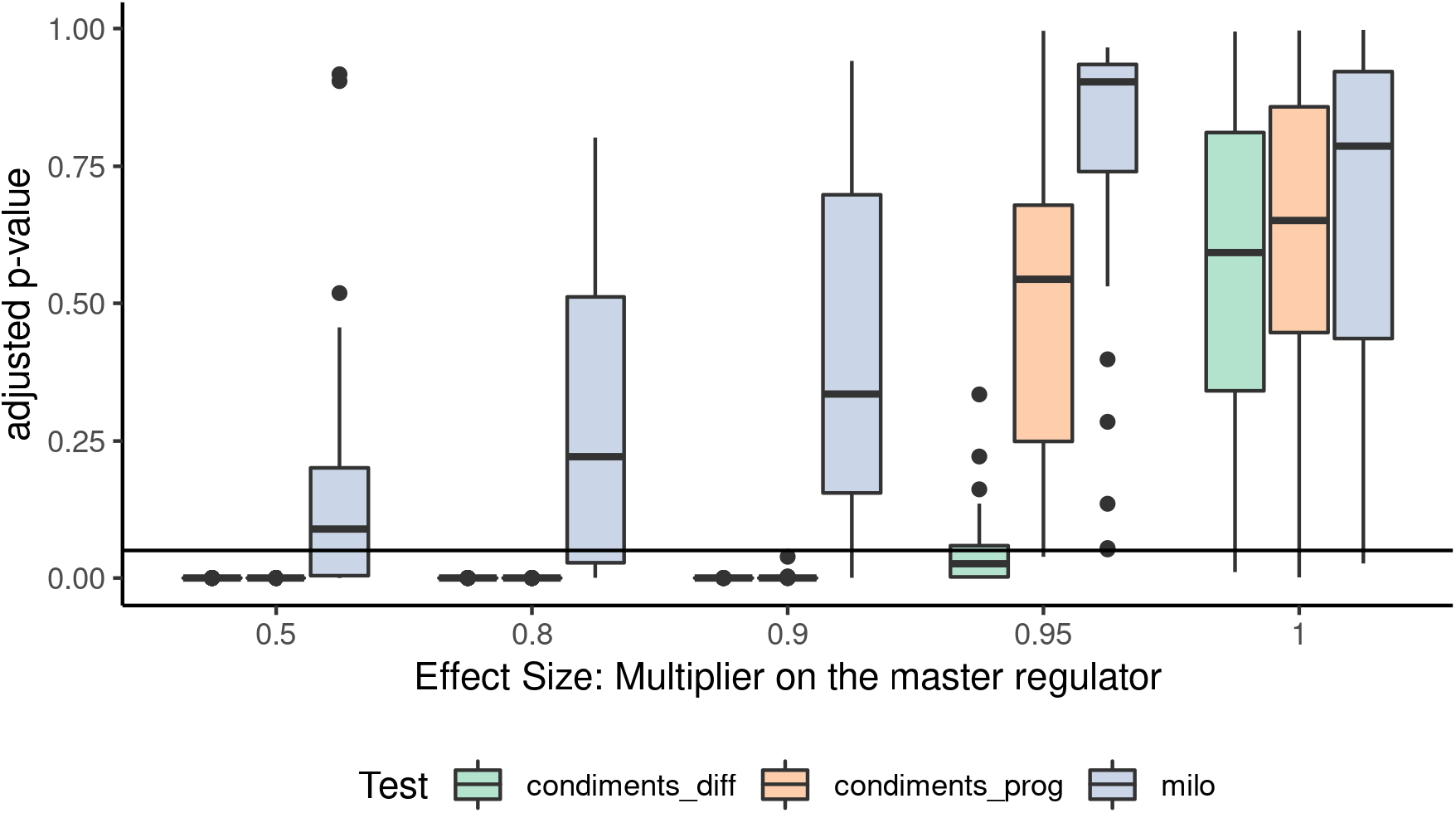
Results on the third type of dataset. For all values of *m* ∈ {.5, .8, .9, .95,1}, we generate null datasets with two lineages and three conditions and we compute the adjusted *p*-values of all tests that can handle 3 conditions. The distributions of *p*-values are then displayed. *m* = 1 is negative (no effect), while *m* < 1 is positive (some effect) with smaller values (toward the left) representing stronger effect.

### S-2.2 TCDD

**Figure S3:**
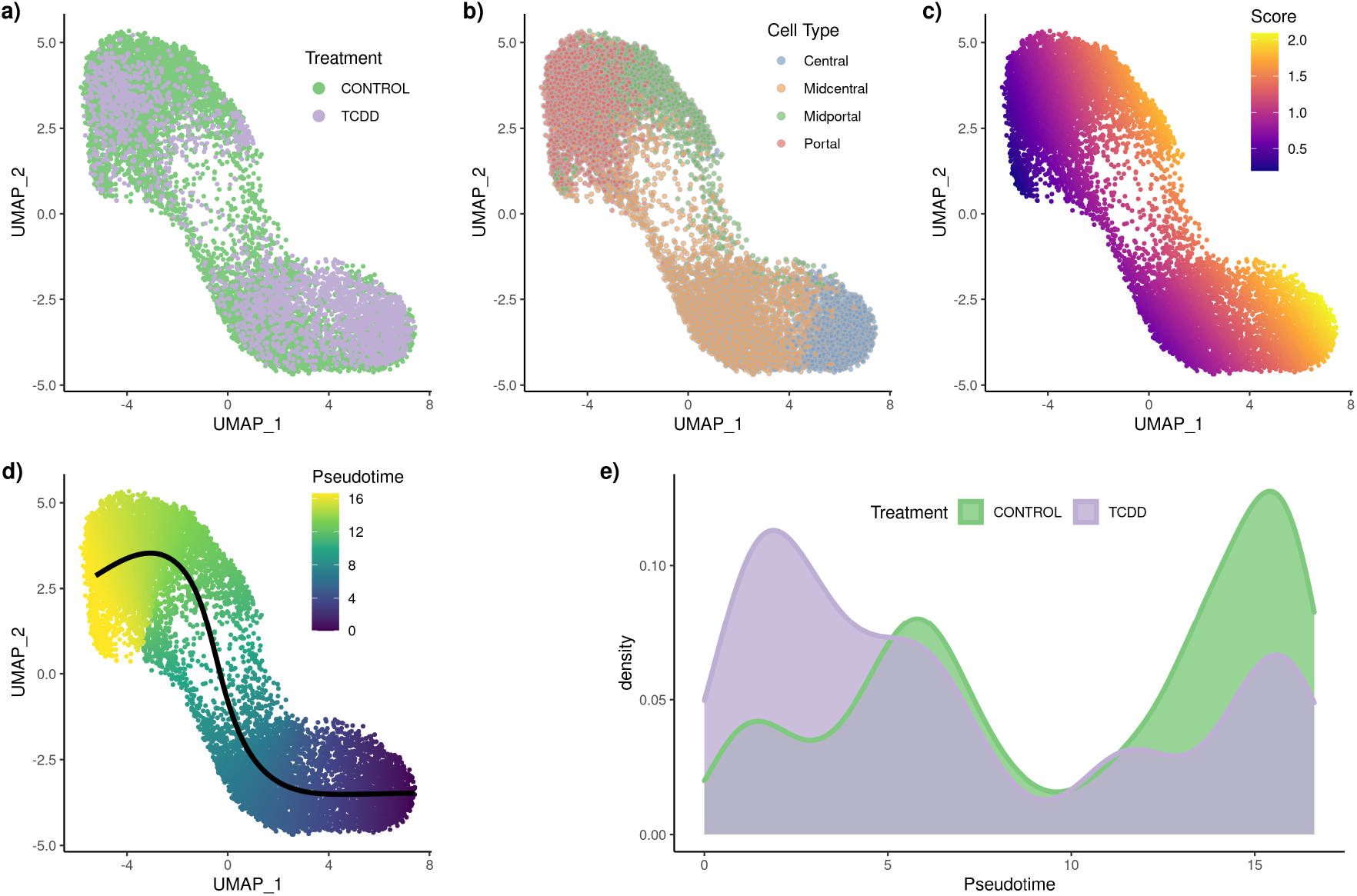
**TCDD** dataset: Differential topology and differential progression. After normalization and projection on a reduced-dimensional space, the cells can be represented, colored either by treatment label **(a.)**, cell type **(b.)**, or batch **(c.)**. Using the treatment label and the reduced-dimensional coordinates, an imbalance score is computed and displayed **(d.)**. The `diffTopoTest` rejects the null and separate trajectories are fitted for each condition **(e.)**. After mapping the lineages, there is also differential progression: the pseudotime distribution along the trajectory are not identical **(f.)** and we indeed reject the null using the `diffProgressionTest`.

**Figure S4:**
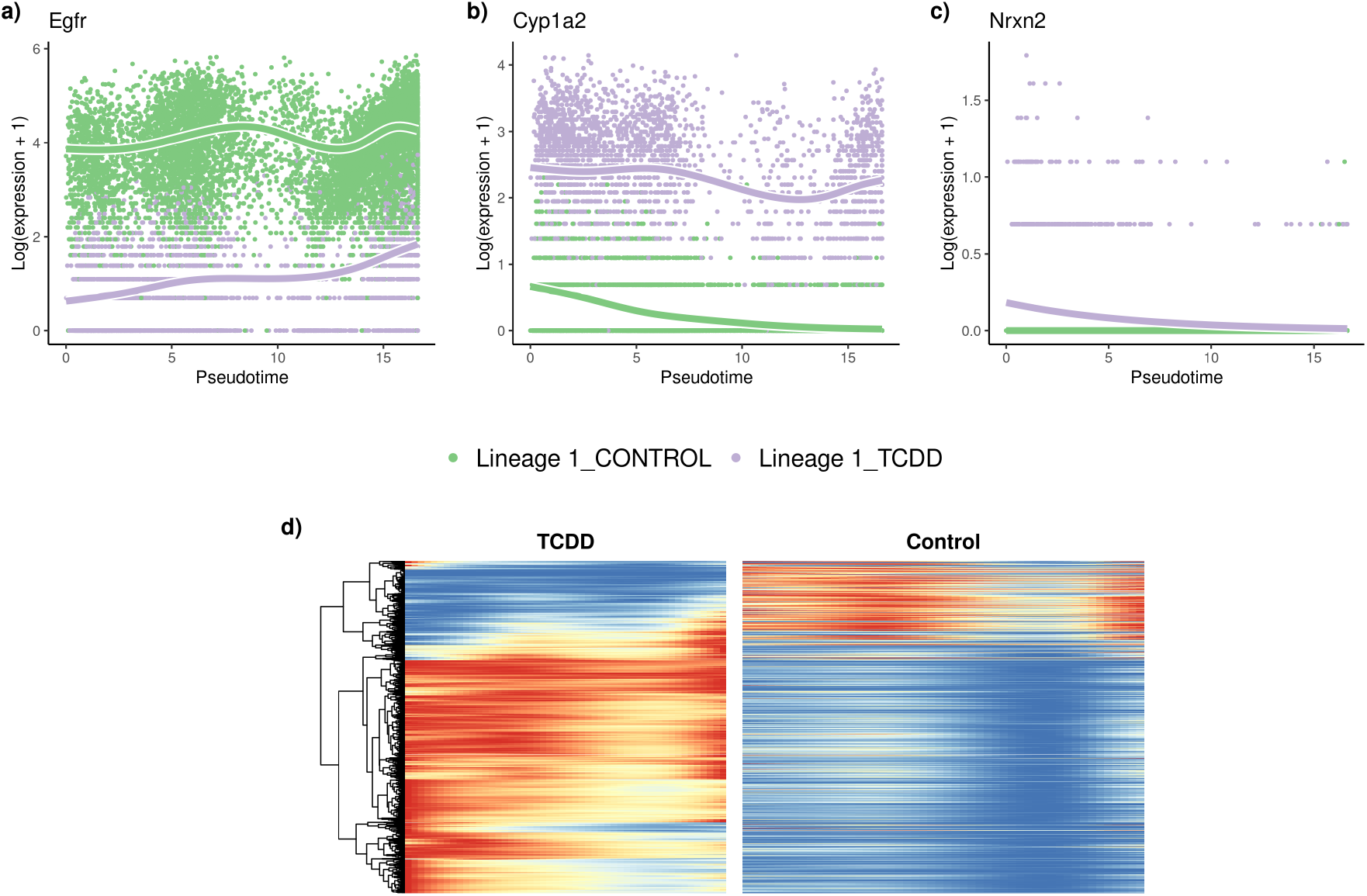
**TCDD** dataset: Differential expression. The `tradeSeq` gene expression model is fitted using the trajectory computed with `slingshot`. Differential expression between conditions is assessed using the `conditionTest` and genes are ranked according to the test statistics. The genes with the highest **(a.)**, second highest **(b.)**, and smallest **(c.)** test statistics are displayed. After adjusting the *p*-values to control the FDR at a nominal level of 5%, we display genes in both conditions using a pseudocolor image **(d.)**.

### S-2.3 KRAS

**Figure S5:**
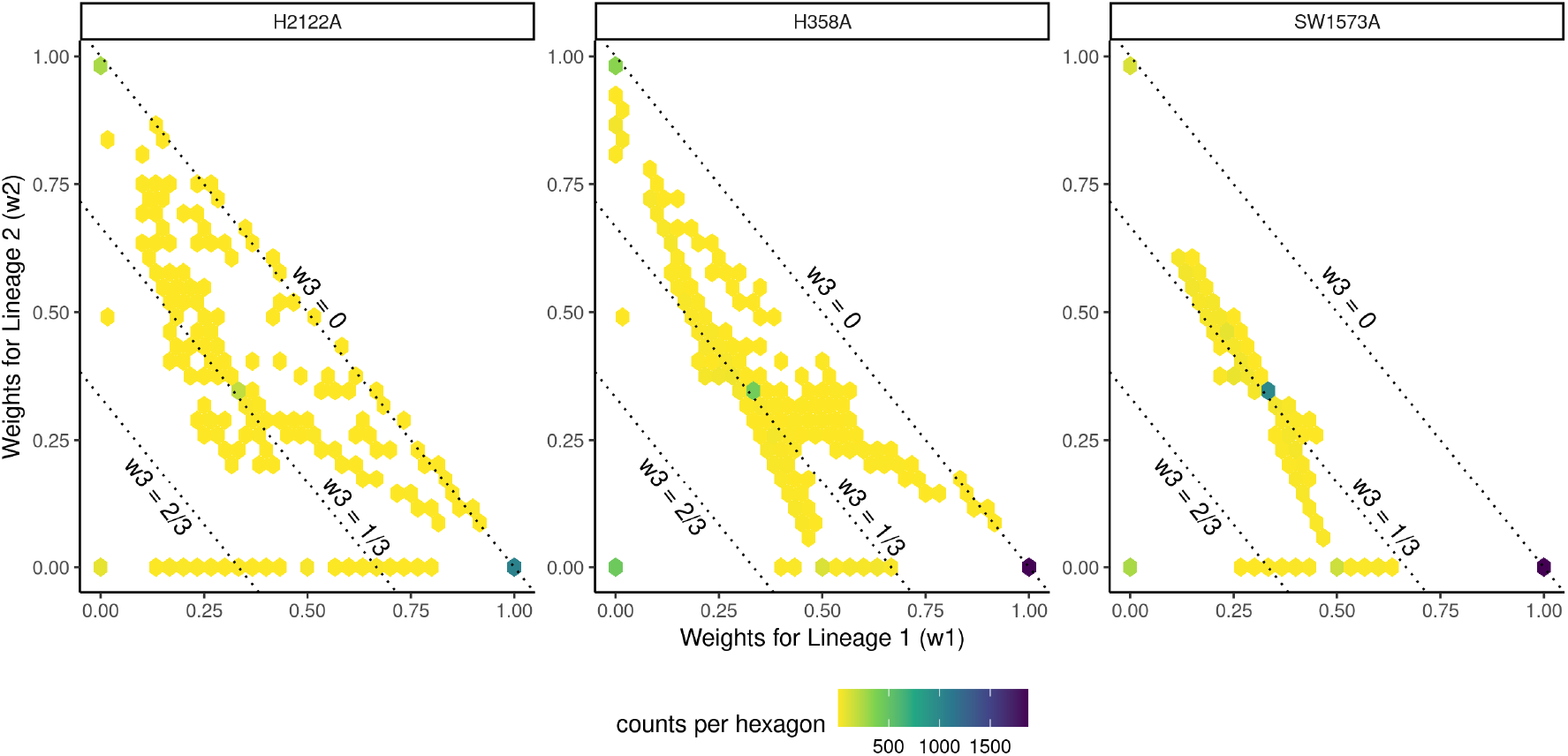
**KRAS** dataset: Differential differentiation.

**Figure S6:**
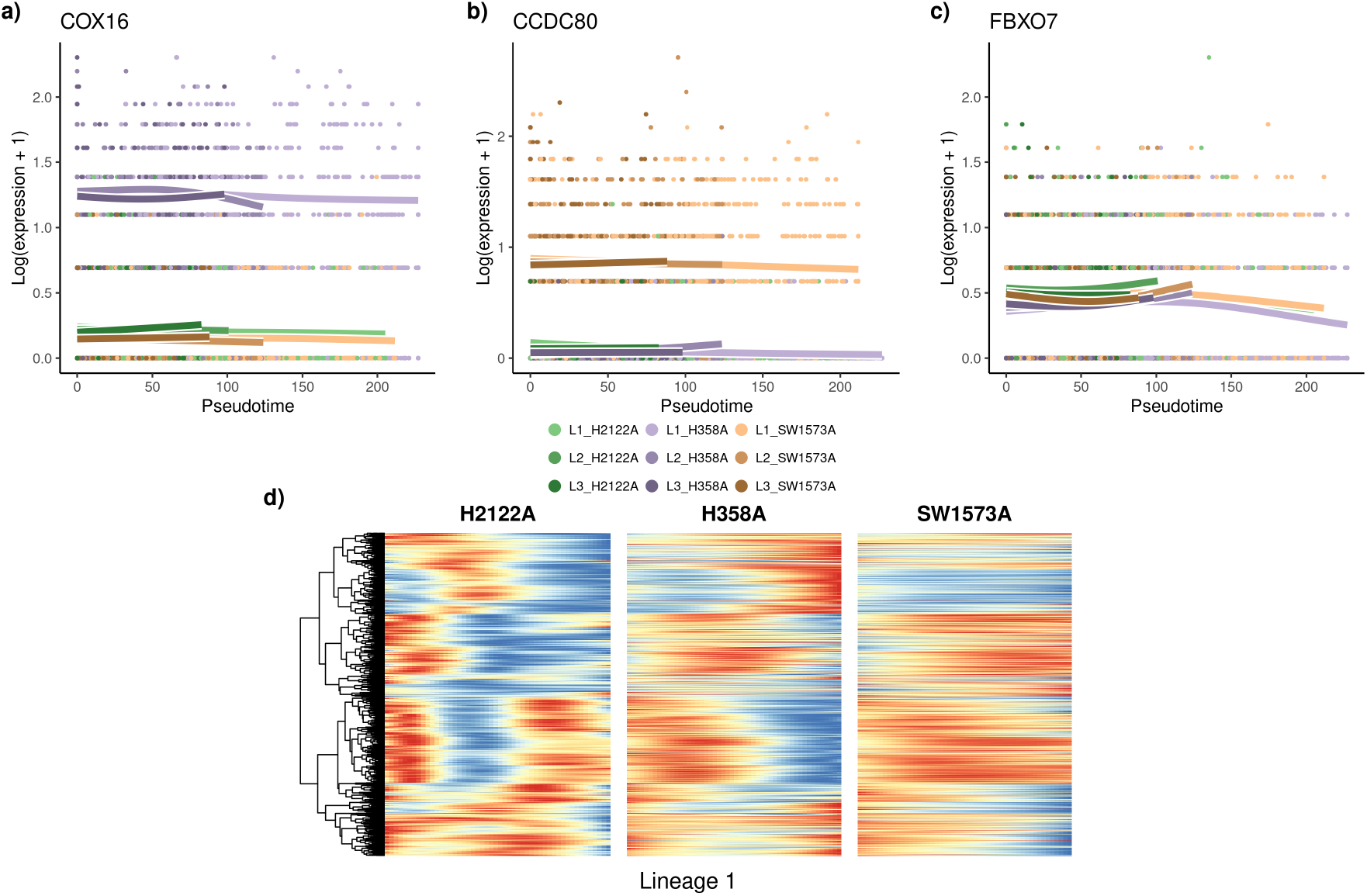
**KRAS** dataset: Differential expression. The `tradeSeq` gene expression model is fitted using the trajectory computed with `slingshot`. Differential expression between conditions is assessed using the `conditionTest` and genes are ranked according to the test statistics. The genes with the highest **(a.)**, second highest **(b.)**, and smallest **(c.)** test statistics are displayed. Focusing on the first lineage, we select all differentially expressed genes in that lineage after adjusting the p-values to control the FDR at a nominal level of 5%. We display the genes for all three conditions using a pseudocolor image **(d.)** along this first lineage.

